# Cooperative *cis*-interactions between ectodomains of TCRαβ and CD3 subunits enable mechanotransduction

**DOI:** 10.1101/2022.04.14.488403

**Authors:** Peiwen Cong, Aswin Natarajan, Zhou Yuan, Chenghao Ge, Stefano Travaglino, Rasool Dezhkam, Saikiran Beesam, Xiang Li, Jiawei Shi, Danielle Grazette, Michelle Krogsgaard, Cheng Zhu

**Affiliations:** Wallace H. Coulter Department of Biomedical Engineering, Georgia Institute of Technology, Atlanta, GA, USA; Petit Institute for Bioengineering and Bioscience, Georgia Institute of Technology, Atlanta, GA, USA; Laura and Isaac Perlmutter Cancer Center and Department of Pathology, New York University Grossman School of Medicine, New York, NY, USA; Georgia W. Woodruff School of Mechanical Engineering, Georgia Institute of Technology, Atlanta, GA, USA

## Abstract

TCR signaling poses a mechanical problem: pMHC binding occurs at the TCRαβ ectodomain head, whereas ITAM phosphorylation occurs on CD3 cytoplasmic tails. Chemistry cannot bridge this > 10 nm gap, so the two events must be coupled through the TCR–CD3 interface, involving conformational allostery and allowing force-based regulation. Although weak ECD *cis*-interactions between TCRαβ and CD3 have been proposed to contribute to this coupling, their kinetics and mechanical competence remain elusive. Here, we quantify TCRαβ–CD3 ECD *cis*-interactions in a pseudo-*cis* configuration by two-dimensional affinity and single-bond force spectroscopy, finding that TCRαβ–CD3γε and TCRαβ–CD3δε bimolecular interactions have low affinity and rapid kinetics, yet form catch bonds. Critically, concurrent engagement of CD3γε and CD3δε generates *cis*-cooperativity, yielding a stronger and longer-lasting CD3γε–TCRαβ–CD3δε trimolecular catch bond than the sum of the two dimeric bonds, with force-stabilized lifetimes matching those of agonist TCR–pMHC *trans*-interaction. Molecular dynamics simulations reveal an expanded, cooperative, and asymmetric contact network, making CD3δε more force-responsive and susceptible to conformational change than CD3γε. Interface mutations do not alter force-free affinities but remodel force-dependent *cis*-bond profiles, leading to an inverse correlation with *trans*-bond profiles and T cell signaling. These results define cooperative ECD *cis*-interaction as a mechanically regulatable allosteric coupling element at the TCR–CD3 junction important to antigen recognition and signal initiation.

## INTRODUCTION

Antigen recognition by the T-cell receptor (TCR) results in T cell activation, leading to proliferation, differentiation, and effector functions (*1*). Signaling is initiated by ligation of peptide-major histocompatibility complex (pMHC) with TCR at its extracellular ligand-binding site, where the antigen information is received, then transmitted across the membrane along the octameric TCR–CD3 complex (TCRαβ associated with CD3γε, CD3δε, and CD3ζζ) to the CD3 cytoplasmic tails (*2*). Concurrently, coreceptor CD8 or CD4 places the Src family kinase Lck close to the exposed CD3 tails to phosphorylate their immunoreceptor tyrosine-based activation motifs (ITAMs) (*3*). Phosphorylated ITAMs recruit Zap70, starting downstream signaling cascades (*1, 4*). Despite our detailed knowledge of the structural components of the TCR signaling machinery (*5, 6*), the mechanism by which pMHC binding at the TCRαβ head leads to phosphorylation of CD3 ITAMs, two events separated by ∼ 15 nm in space, precluding the action of local chemistry, remains unclear.

Two of the TCR triggering models—conformational change and mechanosensor models (*7–9*)— share a common requirement: a physical coupling pathway must link ligand binding to ITAM phosphorylation. Structurally, this pathway must traverse the interface between TCRαβ and CD3s, including ectodomains (ECDs), connecting peptides (CP), the transmembrane (TM) bundle, and cytoplasmic tails. Yet identifying this coupling has been challenging because the TCR faces two seemingly opposing design constraints. On the one hand, the complex must be sufficiently connected to maintain quaternary integrity and transmit information. On the other hand, it must remain sufficiently dynamic to permit state transitions that propagate signals. Resolving how dynamic molecular architecture balances connectivity with mobility is a central obstacle to a unified physical mechanism of TCR triggering.

A likely candidate for this coupling is *cis*-interaction (*i.e.*, lateral interaction between proteins anchored to the same cell surface) between the TCRαβ constant domains (Cα/Cβ) and the CD3 ECDs. Multiple lines of evidence support this notion: the functional sidedness of CD3s (*10*) and the importance of the Cα/Cβ (*11, 12*) indicate that TCR–CD3 *cis*-interactions are crucial for signal transmission. Disrupting these interactions with anti-CD3 antibodies can change how T cells are activated (*13–15*). The membrane-proximal ECD “ring” may act as the starting point for signal propagation and influence downstream elements such as the CPs (*16*), TM bundle (*17, 18*), and cytosolic tails (*2, 19*). Mutations affecting these *cis*-interactions can impact TCR assembly stability (*20, 21*), surface expression (*22*), down-modulation (*23*), and cellular responses (*10, 21, 24–27*). However, two challenges have limited direct mechanistic interpretation. First, ECD interactions appear very weak and transient, raising doubts about their ability to support sustained signal transmission. Second, the subtle coupling may rely on force-dependent dynamic allostery that is sensitive to experimental perturbations, rather than on easily captured large, stable structural changes, as suggested by discrepant results from different experiments: Early cryo-electron microscopy (cryo-EM) structures (*28–31*) and crosslinking experiments (*27*) have shown minimal rearrangements upon ligation, whereas nuclear magnetic resonance (NMR) (*32, 33*) and molecular dynamics (MD) (*33, 34*) studies indicate dynamic allostery in Cα/Cβ and at the CD3 interfaces upon pMHC binding.

More recent cryo-EM structures of the TCR–CD3 complex reconstituted in nanodiscs suggest unliganded TCR–CD3 adopts “compact/closed” conformations (*35*). Upon pMHC engagement, the complex shifts to an “extended/open” architecture, and restricting ECD opening can impair activation (*35*). These findings support the plausibility of activation-linked allosteric transitions, but they do not identify the mechanical mechanism that stabilizes and coordinates this closed-to-open transition across the TCR–CD3 junction, nor do they establish whether the implicated ECD interfaces are mechanically competent to support signal transmitted from TCR–pMHC *trans*-interaction in the presence of force (*1, 6, 18, 36–39*).

Mechanical forces have been suggested to induce conformational allostery (*40–42*), regulate lifetimes of TCR–pMHC *trans*-bonds (*40, 41, 43–46*), and trigger signaling (*43, 47–49*), which, in turn, modulate effector functions (*41, 43, 45, 50, 51*). T cells exert endogenous forces on the TCR–CD3 via engaged pMHC (*52–55*), which must be mediated, at least in part, by TCR–CD3 ECD *cis-*interactions, indicating a possible role for force in regulating these interactions. Thus, mechanical force naturally fits in the allosteric framework, likely as a regulator rather than an alternative to allostery. T cells exert forces on engaged ligands, and mechanical load can bias conformational ensembles by stabilizing specific bound states and reaction pathways. In this view, force is a control parameter that can reveal latent allosteric couplings that are otherwise too fleeting to detect at equilibrium. If ECD *cis*-interactions participate in signal transmission, they must satisfy a pair of stringent mechanical criteria: remaining connected long enough under physiological loads to couple ligand engagement to downstream phosphorylation, yet permitting the local rearrangements and fluctuations required for signal propagation. These seemingly opposing criteria frame our inquiry into a pair of quantitative questions: 1) What are the kinetic and force-dependent properties of the TCRαβ–CD3 ECD *cis*-bonds? 2) Can cooperative assembly transform weak pairwise contacts into a durable, force-stabilized coupling element?

Addressing these questions has been difficult because TCRαβ–CD3 ECD *cis*-interactions are too weak and transient to be measured by three-dimensional (3D) methods (*24, 56–58*) such as SPR (fig. S1, A and B). Moreover, most prior characterizations have been performed without controlling mechanical load, obscuring whether force reshapes the interaction landscape in ways relevant to mechanotransduction. Thus, despite longstanding hypotheses that ECD *cis*-interactions link antigen recognition to CD3 signaling at least partly, direct measurements of their kinetics, cooperativity, and force response have been lacking.

Here, we combine ultra-sensitive biophysical measurements with molecular dynamics (MD) simulations to define the kinetic and mechanical properties of TCRαβ–CD3 ECD *cis*-interactions. Using micropipette adhesion frequency (MAF) assay (*59, 60*) and biomembrane force probe (BFP) thermal fluctuation and force-clamp assays (*43, 61*), we analyzed two-dimensional (2D) binding kinetics of TCRαβ interactions with either CD3γε, CD3δε, or both in a pseudo-*cis* configuration. We found that the bimolecular interactions are low-affinity and short-lived, yet form catch bonds, indicating force-favored bound states. Crucially, co-engagement of CD3γε and CD3δε generates strong synergies in both bond formation and lifetime, resulting in a long-lived cooperative CD3γε– TCRαβ–CD3δε trimolecular catch bond. Complementary atomic-level analyses using conventional MD (CMD) and steered MD (SMD) simulations corroborate cooperative stabilization through an expanded contact network and reveal asymmetric force responses between CD3γε and CD3δε. Finally, targeted mutations at the trimeric interface remodel force-dependent *cis*-bond profiles in a way that inversely correlates with *trans*-bond profiles, TCR mechanosignaling, and T cell activation, yet without altering force-free affinity. Together, these results establish cooperative ECD *cis*-interaction as a mechanically regulatable allosteric coupling element that stabilizes and shapes information transmission across the TCR–CD3 interface.

## RESULTS

### 2D kinetic measurements reveal weak TCRαβ–CD3 *cis*-interactions

To directly measure physical interactions among TCRαβ, CD3γε, and CD3δε (ECDs; hereinafter omitted for simplicity), we utilized the ultra-sensitive MAF assay (*59, 60*) and BFP thermal fluctuation assay (*60, 62*). To compensate for the anticipated low affinities, we coated high densities of hybrid 2B4 TCRαβ (mouse 2B4 TCRαβ variable domains fused with human LC13 TCRαβ constant domains) (*26, 63*) and human CD3γε or CD3δε onto opposing surfaces (Fig. 1A, B). To mimic *cis*-interactions, N-terminal biotinylated TCRαβ was coated on one side (*left*), while C-terminal biotinylated CD3s were coated on the opposing side (*right*), allowing head-to-tail alignment for “pseudo-*cis*” contacts. In the MAF assay (Fig. 1A), one red blood cell (RBC) was driven to cyclically contact the other RBC with a consistent contact area (*A*_c_, in μm^2^) and time (*t*_c_, in s) and separated to detect binding events from visually observed RBC elongations. An adhesion frequency, *P*_a_ (*i.e.*, number of binding events divided by total touches), was enumerated from a sequence of repeated contact cycles. Assuming second-order forward and first-order reverse reactions between monomeric TCRαβ and CD3γε or CD3δε, we previously showed that *P*_a_ = 1 − exp(−⟨*n*⟩), where ⟨*n*⟩ = *m*_r_*m*_l_*A*_c_*K*_a_[1 − exp(−*k*_off_*t*_c_)] is the average number of bonds per contact. Here, *m*_r_ and *m*_l_ (in µm^−2^) are the respective densities of TCRαβ and CD3γε or CD3δε, *K*_a_ (in μm^2^) is 2D affinity, and *k*_off_ (in s^−1^) is off-rate (*59, 60*). When *k*_off_*t*_c_ ≫ 1, exp(−*k*_off_*t*_c_) ≈ 0, the effective 2D affinity *A*_c_*K*_a_ (in μm^4^) can be solved explicitly as *A*_c_*K*_a_ = − ln(1 − *P*_a_) /*m*_r_*m*_l_.

**Fig. 1.**
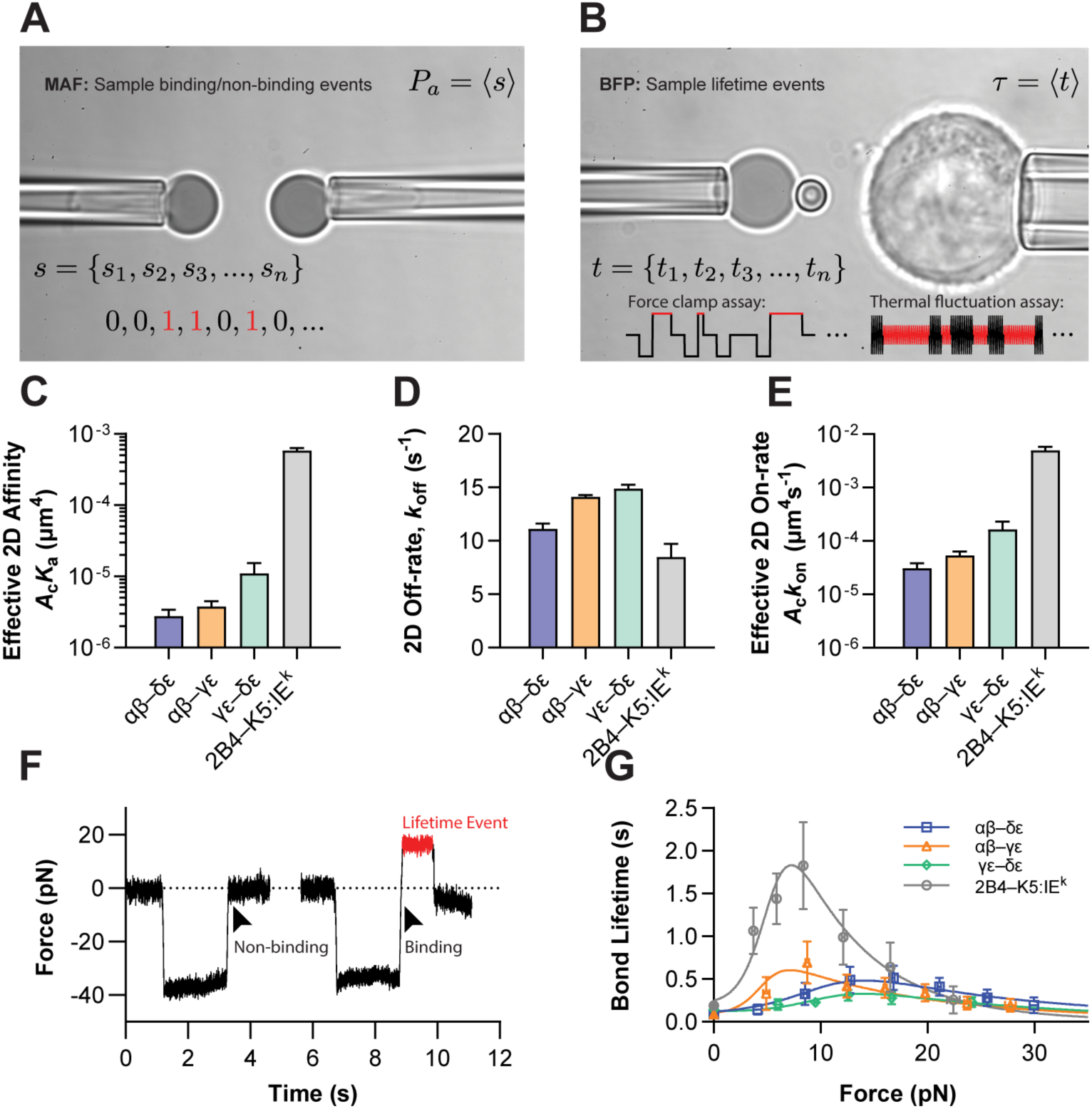
2D kinetic quantifications of TCRαβ–CD3 ECD *cis*-interactions. (**A** and **B**) Photomicrographs of the micropipette adhesion frequency (MAF) assay (A) and the biomembrane force probe (BFP) assay (B), overlaid with legends indicating the measurement types. The two micropipettes held the probe (*left*) and the target (*right*), respectively, with each presenting one of the interacting pair. The target image shown in (B) is a hybridoma cell expressing full TCR–CD3 complexes, but beads coated with purified protein ECDs were also used, depending on the experiment. The target was brought into repeated contact with the probe to enable and interrogate ligand-receptor interactions. The probe, which was either a pressurized RBC (A) or a glass bead attached to the apex of the pressurized RBC (B), served a dual role of ligand presenter and force transducer, detecting the binding events to evaluate adhesion probability (A) and measuring bond lifetime (B), respectively, as indicated in the legends (see Methods). (**C**) Effective 2D affinities (*A*_c_*K*_a_) of pairwise *cis*-interactions between the ECDs of TCRαβ (αβ), CD3δε (δε), and CD3γε (γε) measured by MAF. (**D**) 2D off-rates (*k*_off_) of the indicated *cis*-interactions quantified by the thermal fluctuation assay using BFP. (**E**) Calculated effective 2D on-rates (*A*_c_*k*_on_ = *A*_c_*K*_a_ × *k*_off_) of the indicated *cis*-interactions. Data are presented as mean ± SEM in (C) to (E), with the 2D affinity, off-rate, and on-rate of the *trans*-interaction between cell-surface 2B4 TCR and its cognate pMHC K5:I-E^k^ (*gray*) also plotted in the corresponding panels for comparison. Obtained kinetic parameters are presented in table S1. (**F**) Representative force traces of non-binding *vs.* binding events from the BFP force-clamp assay. The target, either a bead or a cell, was driven to approach the probe (0 pN before contact), pressed against it for 1 s (–35 pN), which resulted in either a non-binding (returning to 0 pN) or a binding event (crossing 0 pN to a positive, tensile force) upon target retraction. The latter trajectory illustrates a lifetime event in which the ligand-receptor bond was clamped with a force of 16 pN for 0.5 s (*red*) until dissociation, indicated by a sudden drop in force to 0 pN. (**G**) Force-dependent average lifetimes of pairwise αβ–δε, αβ– γε, and δε–γε *cis*-bonds (*points*, mean ± SEM of > 30 lifetime events per force bin) at zero force by thermal fluctuation assay (*closed symbols*) and at non-zero forces by force-clamp assay (*open symbols*). The overlaid smooth curves are the global fits of the two-state dissociation model (fig. S2; see Methods). In contrast to *cis*-bonds, the points and curve in *gray* denote the previously published force-lifetime results of the 2B4–K5:I-E^k^ *trans*-interaction (*42*), providing a direct comparison.

The effective 2D affinities of TCRαβ–CD3γε and TCRαβ–CD3δε *cis*-interactions were similar (3.78 and 2.78 × 10^−6^ μm^4^, respectively), which are reliable because they are about an order of magnitude above the detection limit of our current MAF setups (Fig. 1C, *orange* and *blue*) (*46*). For comparison, we measured the TCR–pMHC *trans*-interaction side-by-side using the C-terminal biotinylated K5:I-E^k^-coated RBC to test against the intact TCR–CD3 complex expressed on 2B4 hybridoma cells. The resulting effective 2D affinity was tens to hundreds of folds higher (5.87 × 10^−4^ μm^4^, Fig. 1C, *gray*) even though the hybridoma cell surface is rough and the TCRαβ-bearing RBC is smooth, which could result in one to two orders of magnitude lower effective 2D affinity for the rough than smooth cells (*64*). In addition, using C-terminal biotinylated CD3γε and CD3δε coated on opposing surfaces, we quantified the CD3γε–CD3δε heterotypic interaction (1.10 × 10^−5^ μm^4^, Fig. 1C, *green*), which was higher than the TCR–CD3 *cis*-interactions. Similarly, the effective 2D affinities for the CD3γε–CD3γε, CD3δε–CD3δε, and TCRαβ–TCRαβ homotypic interactions were measured with lower values or were undetectable (fig. S1C).

To further determine the on- and off-rate, we used the BFP thermal fluctuation assay to measure off-rate *k*_off_ (*60, 62*). We attached a functionalized glass bead to the apex of the micropipette-aspirated ultra-soft hypotonic RBC to form the probe on the left (Fig. 1B), whose thermally driven fluctuations were monitored in real time with high spatial-temporal precision (few nanometers and sub-milliseconds) (*65*). The target bead coated with the corresponding binding partner was brought close to the probe to allow for molecular contacts via thermal fluctuations, facilitating bond formation. Upon binding, the receptor-ligand (*i.e.*, either TCRαβ *vs.* CD3, CD3 *vs.* CD3, or pMHC *vs.* TCR–CD3) bond acted as an additional spring in parallel with the RBC spring, dampening the confined Brownian motions (*60, 62*). The sudden reduction in thermal fluctuations indicated bond formation, while the restoration to the original fluctuation amplitudes marked bond dissociation. The durations between events represent bond lifetimes, which were exponentially distributed, and the reciprocal of the mean bond lifetime equals the *k*_off_. The 2D off-rates for TCRαβ–CD3γε, TCRαβ–CD3δε, and CD3γε–CD3δε *cis*-interactions measured in this way were very rapid (> 10 s^−1^) compared to the slower *k*_off_ of the TCR–pMHC *trans-*interaction plotted alongside in Fig. 1D.

These results also validated the 2 s contact time used in our MAF assay, which was sufficiently long to ensure that the measured adhesion frequencies had reached steady state. Finally, the effective 2D on-rates, *A*_c_*k*_on_, were calculated as *A*_c_*k*_on_ = *A*_c_*K*_a_ × *k*_off_ for the corresponding interactions (Fig. 1E). Together, the very low affinities and fast off-rates explain why SPR failed to detect these weak and rapid TCRαβ–CD3 interactions, underscoring the ultra-sensitivity of our 2D assays (*66*).

### Pairwise *cis*-interactions among TCR–CD3 ECDs form bimolecular catch bonds

We (*41–44, 46, 50*) and others (*40, 45, 51, 67, 68*) have shown that mechanical force modulates TCR–pMHC *trans-*interactions and amplifies antigen discrimination (*40, 43, 44*). To test whether force also modulates dissociation of the TCR–CD3 *cis*-interactions, we used the BFP force-clamp assay (*41, 43–45*) to measure the force-dependent lifetimes of TCRαβ–CD3γε, TCRαβ–CD3δε, and CD3γε–CD3δε *cis*-bonds (Fig. 1B). Like in the thermal fluctuation assay, we also used bead-mounted RBCs as ultra-sensitive force transducers to measure molecular interactions in the force-clamp assay. Unlike the thermal fluctuation assay, which positioned the target bead at the null position for bond formation and dissociation at zero force, the force-clamp assay retracted the target by a preset distance to exert a constant tensile force onto the bond. To improve force-measurement stability, we implemented a dual-edge tracking system to monitor the positions of both the BFP micropipette and the probe, allowing us to subtract micropipette drift from the force signals. The target was programmed to move in repeated “approach-touch-retract-hold-return” cycles, generating a force *vs.* time trace (Fig. 1F) that exemplified both non-binding and binding events. The compressive (negative) force indicates bead-target contact during which bond formation can occur. Upon retraction, the compressive force either returns to zero (no binding) or converts to tensile (positive) force (binding). In the latter case, retraction was halted at a given force until bond dissociation, indicated by the sudden return of force to zero. Interestingly, BFP measurements revealed that all three *cis*-interactions formed catch bonds, with mean lifetimes prolonged by low forces (< 7-17 pN, depending on the specific interaction) until being overpowered by higher forces that turned the bond profiles into slip bonds (*i.e.*, mean bond lifetimes shortened by forces; Fig. 1G). These findings demonstrate that TCR–CD3 *cis*-interactions can bear durable force, supporting the notion that information on antigen recognition can be transmitted mechanically through TCRαβ–CD3 interactions.

Notably, these TCRαβ–CD3 *cis*-interactions exhibited shorter bond lifetimes across the entire force range than our previously reported TCRαβ–pMHC *trans*-bonds (replotted from (*42*) in Fig. 1G, *gray*), suggesting that individual TCRαβ–CD3 bimolecular *cis-*interactions alone may not persist long enough to fulfill the proposed mechanical transmission requirements. The 1:1:1 stoichiometry of TCRαβ:CD3γε:CD3δε revealed by the cryo-EM structures (*28, 29, 31*) and our direct kinetic measurements of CD3γε–CD3δε interactions (Fig. 1, C to E, and G, *green*) prompted us to ask: Does TCRαβ bind CD3γε and CD3δε concurrently to form stronger bonds? If so, is this concurrent binding independent or cooperative? If cooperative, how much is the bond stability enhanced, and does this enhancement meet the mechanical transmission requirements?

### CD3δε and CD3γε interact with TCRαβ cooperatively in bond formation and bond lifetime

To test whether CD3δε and CD3γε interact with TCRαβ cooperatively in *cis*, we measured adhesion frequency and bond lifetime between TCRαβ and both CD3s co-presented concurrently. The results were compared against predictions from a non-cooperative binding model using measurements between TCRαβ and the two CD3s presented one at a time (Fig. 2A). Significant deviations from these predictions are taken as evidence for *cis*-cooperativity.

**Fig. 2.**
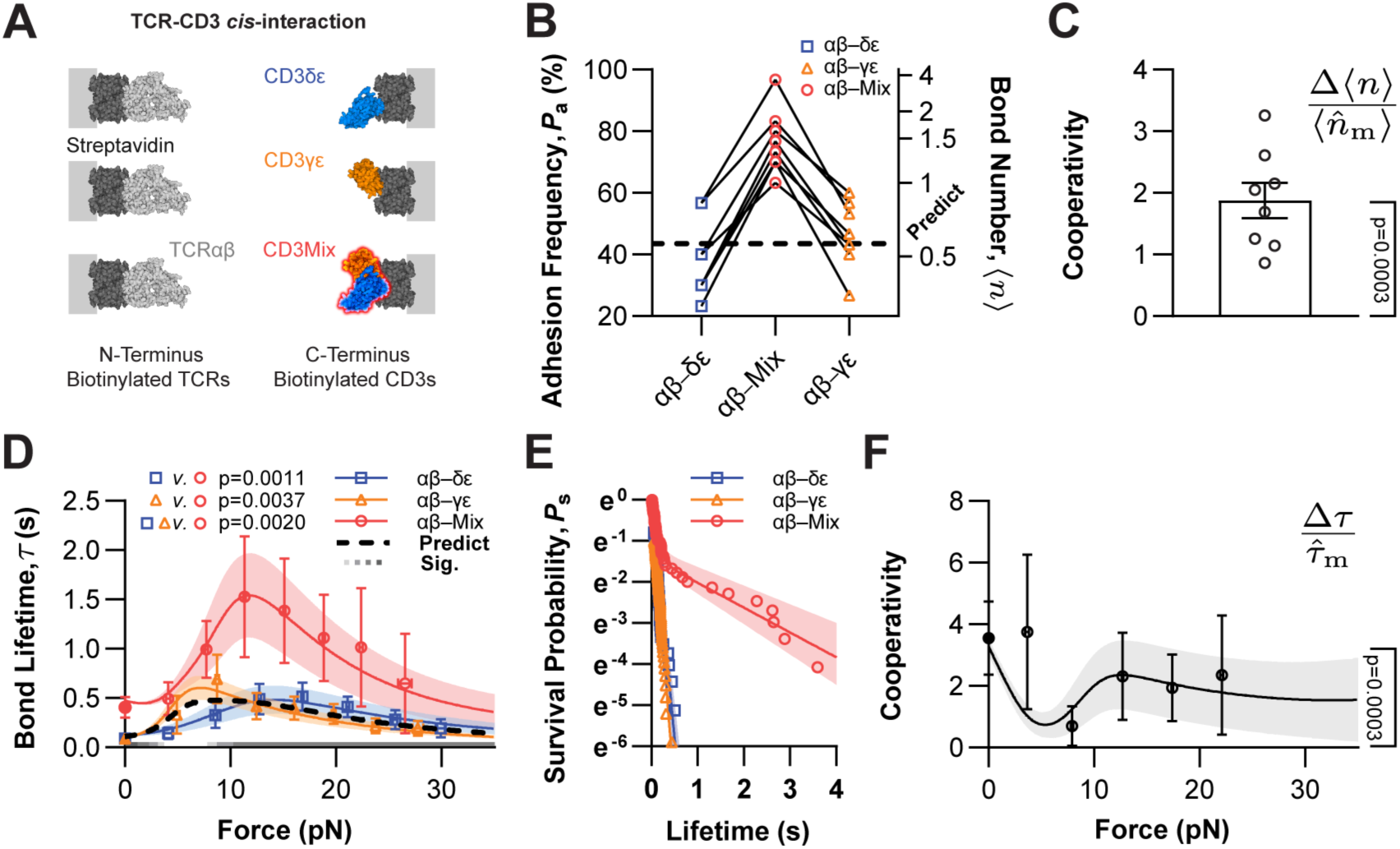
Experimental demonstration of cooperativity in *cis*-interaction of CD3γε, CD3δε, and TCRαβ ECDs. (**A**) Experimental schemes. Purified protein ECDs were captured onto the RBC or bead surfaces through biotin-streptavidin (SA) coupling. N-terminally-biotinylated 2B4 hybrid TCRαβ was coated onto the probe (*left*), while C-terminally-biotinylated human CD3γε (*blue*), CD3δε (*orange*), or a 1:1 mixture of both (CD3m, *red*) was coupled onto the opposing target (*right*). These tail-to-head binding arrangements ensure preferred *cis*-interaction orientations for TCR–CD3 associations. (**B**) Adhesion frequency, *P*_a_ (*left ordinate*), and average bond number, ⟨*n*⟩ = − ln(1 − *P*_a_) (*right ordinate*), were measured by the MAF assay. The same TCRαβ-bearing RBC was tested sequentially against three RBCs bearing, respectively, CD3γε, CD3δε, or CD3m in random order (*points connected by solid lines*). These CD3-bearing RBCs were from the same batch of streptavidinated cells and were incubated with saturating amounts of CD3γε, CD3δε, and CD3m, respectively, to equalize site-density coating. Each probe-target pair was tested for 30 cycles, with a 2-s contact time per cycle. The predicted bond number of dual-ligand RBCs (*dotted line*), ⟨*n̂*_m_⟩, is the sum of the bond numbers of TCRαβ interacting with CD3γε and CD3δε, separately. Thus, ⟨*n̂*_m_⟩ equals the average of the bond numbers measured from the two RBCs bearing single CD3 species because the respective densities of CD3γε and CD3δε on the mixed-ligand RBCs are half of the densities of CD3γε and CD3δε on the single-ligand RBCs under the assumption of concurrent and independent binding (see Methods). This no-cooperativity assumption is invalidated by the data, which clearly demonstrate that the actual measurements greatly outnumber the prediction (note that a log scale is used on the right ordinate). Several metrics were used to quantify cooperativity [(C), (F), and Fig. 3, D and G], which is defined as the difference between the measurement and the prediction, normalized by the expected value assuming no cooperativity. (**C**) Positive 2D binding cooperativity of TCRαβ–CD3m *cis*-interaction. Here, Δ⟨*n*_m_⟩/⟨*n̂*_m_⟩ (*points*) utilized the bond number in (B) as the cooperative metric. No cooperation corresponds to a zero value, whereas a greater-than-zero value indicates 2D binding cooperativity. (**D**) Force-dependent average bond lifetime, *τ*(*f*), for TCRαβ–CD3m *cis*-interactions quantified using BFP. The predicted average bond lifetime of dual-species (*dotted line*), *τ̂*_m_, is the weighted sum of the single-species force-lifetime curves of the αβ–δε and αβ–γε bimolecular bonds (replotted from Fig. 1G for comparison). The weights are fractions of bond formation based on the 2D affinities of single-ligand *cis*-interactions (Fig. 1C) and corresponding amounts (1:1 ratio). The overlaid smooth curves are the global fits of the two-state, two-pathway dissociation models (fig. S2, A to I; see Methods). (**E**) The bond survival probability for the TCRαβ–CD3m *cis*-interactions derived from zero-force individual bond lifetimes (whose average is *τ*(0)) measured from the thermal fluctuation assay. The bond survival probability curves of the αβ–δε and αβ–γε bimolecular bonds are shown for comparison (using pairwise lifetime data in Fig. 1D). (**F**) Positive bond lifetime cooperativity of dual-ligand *cis*-interaction, Δ*τ*/*τ̂*_m_, using the bond lifetime in (D) as the cooperative metric. Again, the no-cooperation case corresponds to a zero value, whereas a value greater-than-zero reveals cooperativity in force-dependent bond lifetime. In (D) and (F), data points are presented as mean ± SEM from > 30 lifetime events per force bin; global-fitting force-lifetime and survival-probability curves (D) to (F) are plotted with means (*solid line*) and standard deviations (*shaded area*). Significance in (C) and (F) was assessed using a one-sample t-test against zero. See Methods for detailed statistical analysis: Pairwise significances ignoring force effect in (D) were obtained using chi-square likelihood ratio tests. Force-dependent significances in (D) were assessed using *z*-tests for mean differences for each force. The gray color bar indicates p-values: ≥ 0.05 (*white*), < 0.05 (*lightest*), < 0.01, < 0.001, to < 0.0001 (*darkest*)

To ensure rigorous comparison of adhesion frequencies, a single TCRαβ-bearing RBC was sequentially brought into contact with an RBC coated with CD3γε, CD3δε, or an equimolar mixture of both (denoted CD3m) in randomized order (Fig. 2A). Strepavidinated RBCs were prepared from the same batch and coated using excess amounts of biotinylated CD3s to ensure saturated binding to SA with equivalent density. Adhesion frequencies were measured over 30 touches per pairing for TCRαβ–CD3γε, TCRαβ–CD3δε, and TCRαβ–CD3m (Fig. 2B, connected by *solid lines*) before switching to a new TCRαβ-bearing RBC (specificity checks of all interactions are shown in fig. S1D). Across eight TCRαβ RBCs tested, the adhesion frequency to CD3m was consistently higher than either subunit alone. To interpret this result, we transformed *P*_a_ into the average number of bonds per contact, ⟨*n*⟩ = − ln(1 − *P*_a_), which is additive for independent, concurrent binding (*69, 70*). Under this non-cooperative assumption, the predicted bond number for CDm is ⟨*n̂*_m_⟩ = 1/2⟨*n*_y**ε**_) + 1/2⟨*n*_δ**ε**_⟩, where ⟨*n*_y**ε**_) and ⟨*n*_δ**ε**_⟩ were measured from respective adhesion frequencies mediated by bimolecular TCRαβ interactions with the single CD3 species as indicated by the subscripts (Fig. 2B, *horizontal dotted line*). Strikingly, the observed ⟨*n*_m_⟩ significantly exceeded the predicted ⟨*n̂*_m_⟩, falsifying the independent binding hypothesis and demonstrating cooperative binding. The degree of cooperativity can be calculated as ⟨*n*_m_⟩/⟨*n̂*_m_⟩ − 1, yielding 170%, substantially higher than zero (Fig. 2C) and confirming pronounced positive *cis*-cooperativity.

Next, we used BFP thermal fluctuation and force-clamp assays to measure the average lifetimes of TCRαβ–CD3m bonds (denoted as *τ*_m_) in the absence and presence of force. The BFP target beads were functionalized similarly to the target RBCs in the MAF assay, except that the surface density of TCRαβ was adjusted to maintain infrequent adhesion events (≤ 20%), ensuring a high probability (*∼* 90%) of single specific bonds (*59*) (fig. S1, E to H). The lifetime scattergram was globally fitted by the previously described two-state, two-pathway dissociation models (fig. S2, A to I) (*46*). TCRαβ–CD3m bonds persisted significantly longer than single-ligand TCRαβ–CD3γε or TCRαβ–CD3δε bonds across all force levels tested (Fig. 2D; single-ligand curves replotted from Fig. 1G). The *τ*_m_ *vs. f* curve peaked at 13 pN with a lifetime of 1.5 s, whereas the *τ*_y**ε**_ and *τ*_δε_ *vs. f* curves peaked at 7 and 15 pN, respectively, with lower values of 0.6 s for both. Notably, the *τ*_m_ bond profile exhibited a more pronounced catching behavior, closely matching that of the TCRαβ– pMHC *trans*-interaction (Fig. 1G, *gray*; peak at 10 pN with a lifetime of 1.6 s). These results indicate that dual ligands are able to provide sufficient *cis-*stability for mechanotransduction, whereas single ligands are not.

To investigate the basis of this prolonged dual-ligand lifetime, we performed survival analysis on bond lifetime distributions (*60, 62, 65*). The survival probability curve of zero-force lifetimes for TCRαβ–CD3m displayed two distinct decay phases, in stark contrast to the log-linear decay observed for single-ligand *cis*-interactions (Fig. 2E). The presence of a slow-decaying subpopulation confirms the existence of the long-lived bond states, bond species, or both. Similar subpopulations of longer-lasting bonds in dual-ligand were also observed in the survival curves of low- and high-force lifetimes (fig. S2, J and K). These longer-lived events likely represent trimolecular CD3δε–TCRαβ–CD3γε bonds that substantially prolong *τ*_m_, and result in positive cooperativity. To quantify the level of cooperativity in terms of bond lifetime, we calculated the average lifetime of TCRαβ–CD3m bonds as the weighted sum of the bond lifetimes of two bimolecular interactions expected from no cooperativity, *τ*_y**ε**_ and *τ*_y**ε**_, by *τ̂*_m_ = *P*_y**ε**_ × *τ*_y**ε**_ + *P*_δ**ε**_ × *τ*_δ**ε**_ (Fig. 2D, *dotted curve*), where the bond formation possibilities, *P*_y**ε**_ and *P*_δ**ε**_, equal the respective fractions of TCRαβ–CD3γε and TCRαβ–CD3δε bond numbers, ⟨*n*_y**ε**_)/(⟨*n*_y**ε**_) + ⟨*n*_δ**ε**_⟩) and ⟨*n*_δ**ε**_⟩/(⟨*n*_y**ε**_) + ⟨*n*_δ**ε**_⟩). The assumption of independent binding predicts *τ*_m_ = *τ̂*_m_, but the cooperative analysis, Δ*τ*/*τ̂*_m_ = (*τ*_m_ − *τ̂*_m_)/*τ̂*_m_, revealed 74-328% increments in the CD3γε– TCRαβ–CD3δε trimolecular *cis*-bond lifetime over the force range of 0-30 pN (Fig. 2F), highlighting the strong force-enhanced positive cooperativity.

### MD simulations corroborate the cooperative interaction of CD3γε and CD3δε with TCRαβ

To corroborate the experimentally observed positive *cis*-cooperativity, we performed force-free CMD simulations (*71*) on TCRαβ–CD3γε, TCRαβ–CD3δε, and CD3γε–TCRαβ–CD3δε complexes. The initial full-atomic models of TCR–CD3 ECDs, including the trimeric and dimeric structures (Fig. 3A and fig. S3A), were extracted from the cryo-EM structure (PDB: 6JXR) (*28*). All equilibrated complexes remained bound after 200 ns CMD simulations. We analyzed their binding interfaces in terms of intermolecular subunit contact areas (Fig. 3B and fig. S3B) and binding free energies (Fig. 3C and fig. S3C), which were inversely correlated. Using cooperative analyses similar to Fig. 2, C and F, we found these two measures also exhibited positive *cis*-cooperativity (Fig. 3D). Because binding affinity, *K*_a_, is related to free energy change, Δ*E* by *K*_a_ ∝ exp(−Δ*E*/*k*_B_*T*), where *k*_B_ is the Boltzmann constant and *T* is the absolute temperature, these simulations provide independent verification of cooperative 2D affinity measurements.

**Fig. 3.**
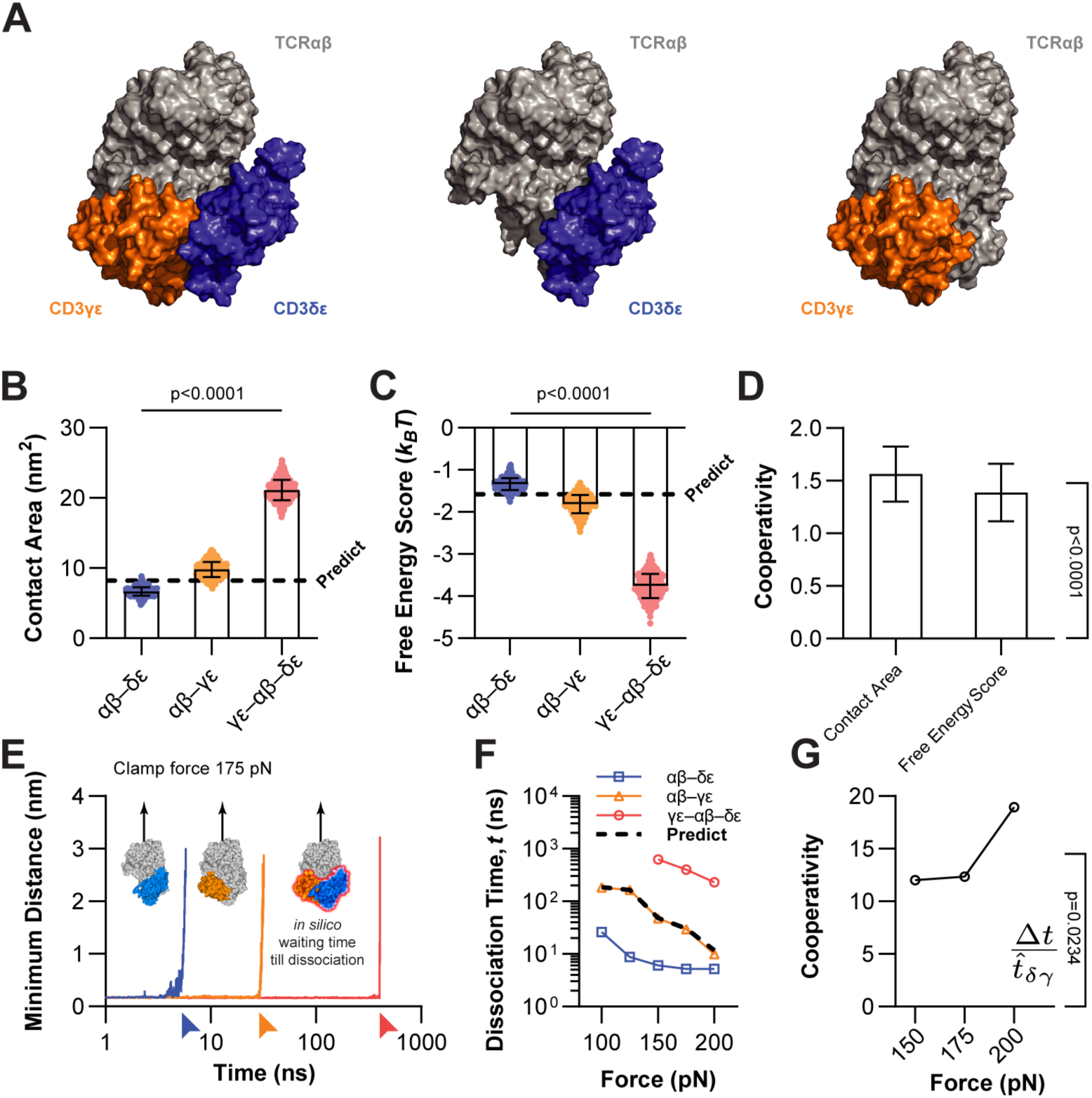
Computational demonstration of cooperativity in *cis*-interaction of CD3γε, CD3δε, and TCRαβ ECDs. (**A**) Initial structures for MD simulations. The ECD of CD3γε–TCRαβ–CD3δε trimolecular complex (*left*) was modeled based on the published cryo-EM structure (*28*), 6JXR. The two TCRαβ–CD3δε (*middle*) and TCRαβ–CD3γε (*right*) bimolecular structures were obtained by removing CD3γε and CD3δε from the trimer, respectively. (**B** and **C**) Contact areas and interaction energies for the dimers and trimer from CMD simulations. Mean ± SD of contact areas (B), defined by solvent accessible surface areas (*108*), and interaction energies (C), quantified using empirical free energy scores (*109*), were calculated from the TCRαβ–CD3δε (*blue*), TCRαβ–CD3γε (*orange*), and CD3γε–TCRαβ–CD3δε (*red*) structures (fig. S3, B and C). Statistics were obtained by comparing any two groups using unpaired Student’s t-tests. (**D**) Cooperative metrics of the δε–αβ– γε trimeric complex in terms of contact areas (*left*) and free energy scores (*right*). The cooperative analyses were conducted similarly to that in Fig. 2C, but were based on contact areas and interaction energies in (B) and (C) rather than on bond numbers. (**E**) The minimum distances between TCRαβ and corresponding CD3δε (*blue*), CD3γε (*orange*), or both (*red*) *vs.* simulation time, calculated from representative trajectories of SMD-simulated TCR–CD3 dissociation. Initial molecular constructs of αβ–δε and αβ–γε dimeric assemblies and of δε–αβ–γε trimeric complex are indicated, where TCRαβ N-termini were pulled away from corresponding C-terminal anchored CD3s during a 175-pN force-clamping SMD simulation (*insets*). The sudden rise of the trajectory from the baseline signifies the time point of dissociation (*arrows*). (**F**) *In silico* times required for the complex to dissociate, *t*(*f*), under constant forces of 100-200 pN from SMD simulations. The predicted time-to-dissociation of the γε–αβ–δε trimer (*dotted line*), *t̂*_δy_, is the TCR dissociation time from both δε and γε, under the assumption that the two bimolecular bonds are in parallel, and their dissociations are independent and concurrent (see Methods). (**G**) Cooperative metric Δ*t*/*t̂*_δy_ *vs.* force of the δε–αβ–γε trimeric complex. The cooperative analyses were conducted similarly to that in Fig. 2F but were based on the SMD simulated time-to-dissociation in (F) rather than on experimentally measured bond lifetimes. Significance was assessed using one-sample t-tests against zero in (D) and (G).

We next simulated force-induced dissociation of TCRαβ–CD3 complexes using force-clamp SMD simulations (*72*). A pulling force was applied on the TCRαβ N-terminus, while the C-termini of CD3γε, CD3δε, or both were anchored (Fig. 3E, *insets* and fig. S4A). To complete simulation within accessible computational timescales, we used relatively high clamping forces (100-200 pN) to accelerate dissociations, which were detected as sudden increases in the minimum distances between TCRαβ and either or both CD3s (Fig. 3E, *curves* and fig. S4B). The times to dissociate ranked as TCRαβ–CD3δε < TCRαβ–CD3γε < CD3γε–TCRαβ–CD3δε. Simulated force effects on time-to-dissociation (Fig. 3F) were consistent with experimental lifetime trends, following expected slip behaviors beyond 15 pN (Fig. 2D). The trimolecular CD3γε–TCRαβ–CD3δε bonds dissociated at 626, 404, and 233 ns under 150, 175, and 200 pN, respectively, which persisted at least an order of magnitude longer than the two dimeric bonds. To quantify cooperativity, we calculated Δ*t*/*t̂*_δy_, which compares observed times to dissociation, *t*_δy_, to the predicted times to dissociation for the non-cooperative trimolecular complex *t̂*_δy_ = *t*_y**ε**_ + *t*_δ**ε**_ − *t*_y**ε**_*t*_δ**ε**_/(*t*_y**ε**_ + *t*_δ**ε**_), calculated using the TCRαβ–CD3γε and TCRαβ–CD3δε dissociation times, *t*_y**ε**_ and *t*_δ**ε**_. Observed dissociation times exceeded this prediction by from 1,200% at 150 pN to 1,900% at 200 pN, demonstrating strong *cis*-cooperativity and mechanical reinforcement (Fig. 3G). Likewise, force-ramp SMD simulations on partial and full TCR–CD3 assemblies yielded rupture forces in the same order, TCRαβ–CD3δε < TCRαβ–CD3γε < CD3γε–TCRαβ–CD3δε (Fig. 4, C and D), further confirming the higher stability of the trimolecular complex. Together, complementary experimental and computational results demonstrate positive *cis*-cooperativity between CD3δε and CD3γε in their co-binding to TCRαβ.

**Fig. 4.**
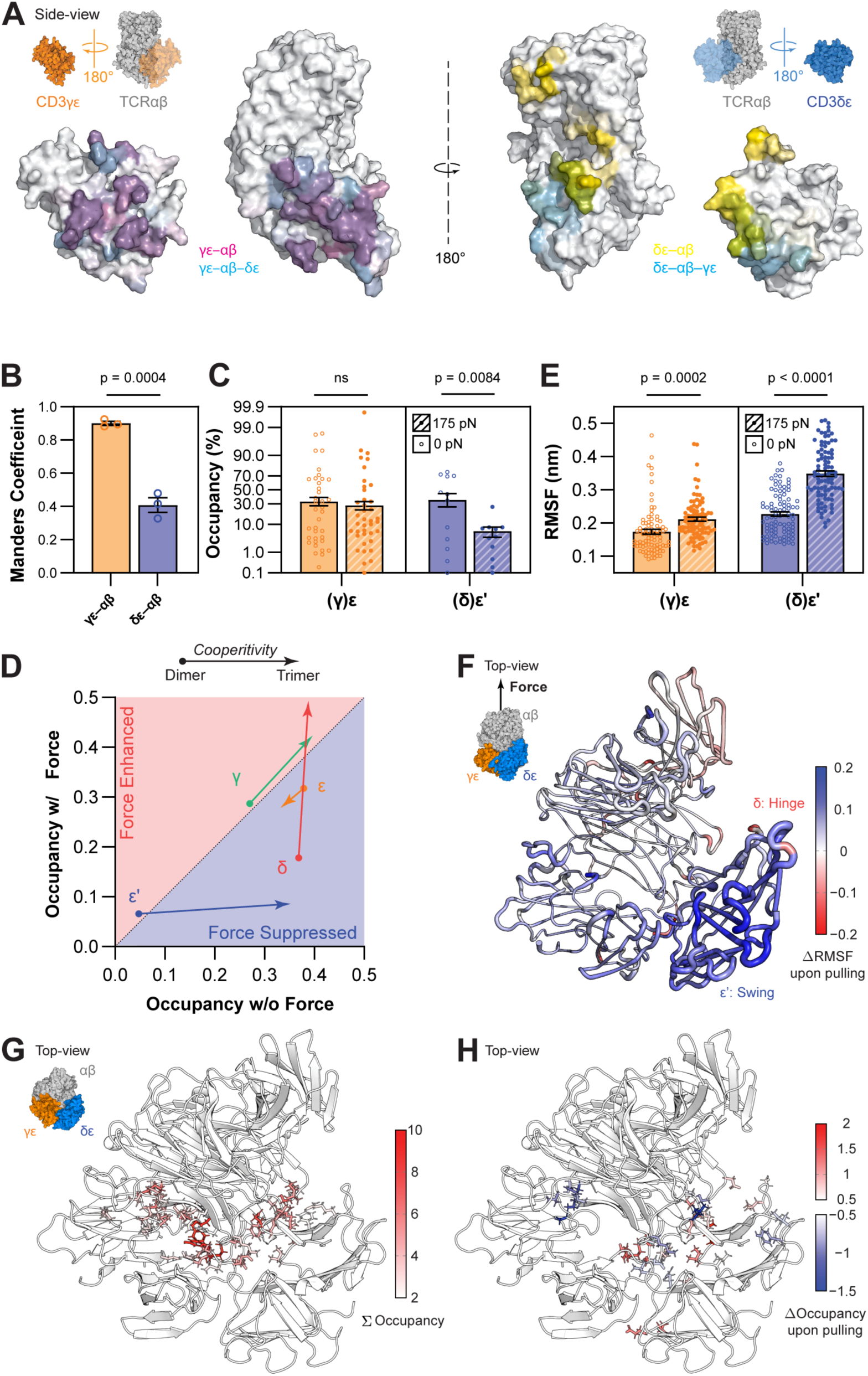
Structural basis of force-dependent cooperative TCR–CD3 ECD *cis*-interactions. (**A**) Dynamic “footprints” of αβ–γε (*left*) and αβ–δε (*right*) interfaces are defined by the CMD-derived occupancies, which were projected onto the light gray space-filling representations of TCR–CD3 ECDs. The *insets* depict the placements and orientations of γε (*orange*) and αβ–δε (*gray*), αβ–γε (*gray*) and δε (*blue*) from the far left to the far right: two sets of side-view images mirroring each other to allow CD3s to face outward. The respective γε (*left*) and δε (*right*) binding interfaces with αβ are exposed by flipping the CD3s 180° while keeping the rest fixed (before *transparent*; after *opaque*). The pseudo-color maps are coded for interface footprints in the trimer (*cyan*; only the unobscured areas are visible) as well as in dimers with γε (*magenta*) and δε (*yellow*), with intensity levels matching the occupancy values, ranging from 0 (white) to 100% (fully saturated). When comparing trimer *vs.* dimer, the unobscured *cyan* areas relative to the *magenta* and *yellow* areas represent the increased footprints of αβ with γε (*left two* panels) or δε (*right two* panels) from dimer structures upon trimer assembly by incorporating the missing CD3. When comparing γε *vs.* δε, the magenta area is larger than the yellow area, indicating a more stable αβ– γε than αβ–δε dimer. Moreover, the overlapping areas are smaller on the right (*cyan* + *magenta*) than on the left (*cyan* + *yellow*), both in absolute values and percentages, indicating smaller cooperative effects due to adding δε to αβ–γε than adding γε to αβ–δε in the trimer and demonstrating asymmetry with less contact adjustment of γε than δε after trimer assembly. (**B**) Quantification of colocalization of occupancies between trimer and dimers by Manders’ overlap coefficient, defined by 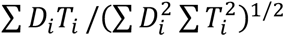 where *i* denotes the residue index of αβ–γε (*orange*) and αβ–δε (*blue*); *D_i_* and *T_i_* denote the *i*^th^ occupancy values of dimer and trimer, respectively, before and after *cis*-cooperative trimer assembly with the missing CD3. A Student t-test was used to assess significance based on three independent simulation repeats in (B). (**C**) Mean ± SEM occupancies calculated from individual residues with non-zero values (*points*) of ε (*left*) or ε’ (*right*) chains interacting with αβ calculated from CMD (0 pN, *open bars*) or SMD (175 pN, *hatched bars*) simulated γε–αβ–δε trimeric structure. Here, γ-associated ε-chain (ε) and δ-associated ε-chain (ε’) are distinguished using the prime superscript. (**D**) Mean occupancies of γ (*green*), ε (*orange*), δ (*red*), or ε’ (*blue*) chains interacting with αβ quantified by averaging non-zero occupancies calculated from CMD (without force, *x*-axis) or SMD (with force, *y*-axis) simulated αβ–γε and αβ–δε dimer structures (*arrow starting points*) or γε–αβ–δε trimer structure (*arrow ending points*). (**E**) Fluctuations of two ε-chains under 0 and 175 pN force, quantified by mean ± SEM of residue RMSFs of ε (*left*) or ε’ (*right*) chains calculated from CMD (0 pN, *open bars*) or SMD (175 pN, *hatched bars*) simulated γε–αβ–δε trimeric structure. Statistical significance levels were assessed by Welch’s t-test assuming non-equivalent SD in (C) and (E). (**F**) Change of flexibilities in the TCR–CD3 complex upon pulling. The *inset* shows the top view of the γε–αβ–δε ECD structure under a 175 pN force. Differential RMSF profiles between SMD and CMD (*blue* for enhanced and *red* for suppressed fluctuations) are mapped onto the ribbon view of trimeric ECD. The residues’ absolute ΔRMSF values are represented by the thickness of the protein backbone. (**G** and **H**) Critical interacting residues mapped onto the γε–αβ–δε interfaces showing the strong-interacting (G) and the force-sensitive (H) contacts. The *inset* in (G) is the top view of the γε–αβ–δε ECD structure. The same top views of the full ECD assembly are shown in ribbon representation, where tightly interacting residues are highlighted by sticks and colors. Only strong-interacting contact residues are shown in (G) using side chains colored from *white* to *red*, based on their ΣOccupancy level. The force-sensitive residues with significant occupancy differences upon applying a 175 pN clamping force are shown in (H), with side chains colored from *white* to *red* for increases and from *white* to *blue* for decreases in their ΔOccupancy level (see Methods).

### Structural basis of *cis*-cooperativity quantified by atomic-level contacts

To elucidate the structural basis of *cis*-cooperativity, we analyzed interface contacts within the TCRαβ–CD3γε, TCRαβ–CD3δε, and CD3γε–CD3δε dimers and compared them with those within the CD3γε–TCRαβ–CD3δε trimer. The contact area sizes were preserved from the dimer to trimer interfaces, with only modest alteration upon incorporation of the third subunit. Because CD3γε and CD3δε bind TCRαβ orthogonally with minimal overlap, their combined contact area in the trimer was only slightly smaller than the sum of dimer contact areas. This slight steric loss was more than offset by the gain in CD3γε–CD3δε contact area upon trimer formation, which substantially increased total contact area and lowered binding free energy (Fig. 3, B and C), consistent with *cis*-cooperativity. Similarly, we analyzed hydrogen (H) bonds within the TCRαβ–CD3γε, TCRαβ–CD3δε, and CD3γε–CD3δε dimers and compared them with those within the CD3γε–TCRαβ–CD3δε trimer, finding substantially more H-bonds in the trimer than in the dimers at the start and in the middle, but not near the end of SMD simulations immediately before dissociation (fig. S4E). Similar cooperativity calculations found > 100 and 180% more H-bonds in the trimer without and with force, respectively (fig. S4F).

We further quantified the dynamic interfacial contact of each residue to its binding partner using occupancy, defined as the fraction of simulation time during which the center of mass of that residue remains ≤ 4 Å of any atom of its *cis*-interacting ECDs, hence ranging from 0-100%. To reveal dynamic binding footprints, we mapped these occupancy levels onto the molecular surfaces, with color intensities proportional to occupancy percentages (Fig. 4A). To expose the respective TCRαβ–CD3γε and TCRαβ–CD3δε interfaces, a pair of the same trimer with different views forms two mirror images, where the CD3γε (Fig. 4A, *left*) and CD3δε (Fig. 4A, *right*) are rotated outward away from the fixed TCRαβ. Differential color coding is used to indicate CMD-derived occupancy levels on interfaces of CD3γε–TCRαβ–CD3δε trimer (*cyan*), TCRαβ–CD3γε dimer (*magenta*, after CD3δε removal), and TCRαβ–CDδε dimer (*yellow*, after CD3γε removal), respectively. Overlay of dual-color maps (*bluish* overlap for TCRαβ–CD3γε; *greenish* overlap for TCRαβ–CD3δε) highlights conserved interactions. These analyses reveal shifts in residue contacts from the respective dimeric structures to the trimeric structure in the absence of force. Interestingly, the TCRαβ–CD3γε interface is nearly unchanged in the presence of CD3δε (Fig. 4A, *left*, *magenta vs. cyan*), whereas the TCRαβ–CD3δε interface shows a marked reorientation in the presence of CD3γε (Fig. 4A, *right*, *yellow vs. cyan*). This was consistently observed in all three independent CMD runs.

To quantify occupancy footprint colocalization between dimer and trimer states, we calculated Manders’ overlap coefficients (*73*), finding significantly greater values for CD3γε than CD3δε (Fig. 4B). The higher Manders’ coefficient for γε than δε reveals a higher level of overlapping contact interfaces between the trimer and dimer of TCRαβ with CD3γε than CD3δε, which negatively correlates with the level of contact adjustments going from dimer to trimer after adding the missing CD3. This indicates that, structurally, the TCRαβ–CD3γε interface remains unchanged with or without CD3δε, whereas the TCRαβ–CD3δε interface must reshape to accommodate CD3γε incorporation during trimeric assembly. This asymmetric TCRαβ with two CD3s interactions may result in unequal contributions to *cis*-cooperativity.

### Asymmetric responses of CD3ε under *cis*-cooperativity and mechanostransmission

To elucidate the roles of asymmetry of CD3s in their cooperative binding with and force transmission from TCRαβ, we further investigated the differential dynamic responses of CD3γε *vs.* CD3δε using occupancy and fluctuation analyses. We first compared the occupancies determined using SMD simulations in the presence of force with values determined in the previous section using CMD simulations for every single α, β, γ, ε, δ, and ε’ chain (note the omission of TCR and CD3 designations for simplicity). Hereinafter, we use ε’ to denote the ε chain that forms the heterodimer with the δ chain to distinguish it from the ε chain that forms the heterodimer with the γ chain, despite their identical sequences. Notwithstanding the average occupancies of ε’ and ε chains in the trimer are similar in the absence of force, ε’ has far fewer contact residues with αβ than ε both in the absence and presence force, indicating asymmetry (Fig. 4C). Interestingly, under pulling force applied to the TCRαβ, both the number of residues with non-zero values and the domain average of ε’ occupancies decline sharply, whereas those of ε remains stable (Fig. 4C). These data are consistent with the longer lifetime and greater rupture force for TCRαβ–CD3γε than TCRαβ–CD3δε bonds observed in the SMD studies (Fig. 3, E to F, and fig. S4, B to D), suggesting that ε’ is more sensitive to force, whereas ε provides more mechanical support for *cis*-cooperativity.

To expand the ε and ε’ occupancy data (Fig. 4C), we similarly calculated mean occupancies of αβ residues contacting γ and δ domains from CMD and SMD simulated γε–αβ–δε trimer. We also performed parallel occupancy calculations using simulated αβ–δε and αβ–γε dimers for all CD3 chains. We then plotted these paired occupancies as points using values obtained from CMD (without force) and SMD (with force) as *x*- and *y*-coordinates and drew vectors connecting any two points for the same CD3 chains with the start- and end-point coordinates calculated from simulated dimeric and trimeric structures, respectively (Fig. 4D). The magnitude of a vector represents the amount of occupancy change for a single CD3 chain (γ, ε, δ, or ε’) from before (as αβ–γε or αβ–δε dimer) to after (as γε–αβ–δε trimer) incorporating the missing CD3 heterodimer (δε or γε). Remarkably, the δ (*red*) and ε’ (*blue*) vectors were much larger in magnitude than the γ (*green*) and ε (*orange*) vectors, revealing much larger cooperativity-induced occupancy changes in the δ and ε’ than in the γ and ε chains. This quantitatively depicts the strength of larger non-overlapping trimer and dimer footprints (*cyan areas*) of δε (Fig. 4A, *right*) than γε (Fig. 4A, *left*), re-emphasizing that, to form a γε–αβ–δε trimer via cooperative *cis*-interactions among ECDs, it requires more contact adaptation from incorporating γε into the αβ–δε dimer than incorporating δε into the αβ–γε dimer.

The directions of these vectors indicate whether the changes are positive (increasing) or negative (decreasing). Their *x*- and *y*-components represent the changes occurring in the absence and presence of a 175 pN force, respectively. In Fig. 4D, points below (*light blue*) or above (*light red*) the diagonal line indicate domain contacts of αβ with CD3 chains that are suppressed or enhanced by force, respectively. Interestingly, δ switches from a force-suppressed to a slightly force-enhanced state (moving from below to above the diagonal line), while ε’ redirects from slightly force-enhanced to force-suppressed (moving from above to below the diagonal line); by comparison, hardly any changes of force-induced contacts and switches of their force-enhanced (γ) or force-suppressed (ε) states are seen in either γ or ε chain (Fig. 4D). These results reveal that δε is more force-sensitive and γε is more mechanically stable, indicating their different roles in transmitting force across the interfaces between TCRαβ and the two CD3 ECDs.

In addition to occupancy analysis, which measures contact stability, we analyzed root-mean-square fluctuations (RMSF) in the absence and presence of a 175 pN clamp force, which reflect residue flexibility. By averaging across the entire chain, we assessed mobility changes in different chains before and after applying force, revealing another layer of mechano-responsiveness in the trimer. ε’ shows a larger mean RMSF than ε (Fig. 4E), and force-enhances RMSF in both ε’ and ε residues, but much more enhancements for ε’ than ε (Fig. 4E). These results invert the trends of the occupancy data (Fig. 4C), showing a negative correlation expected intuitively. The differences in RMSF values calculated in the presence and absence of force (ΔRMSF) are projected onto the TCR–CD3 quaternary ECD structure, where force-enhanced (*blue*) and force-suppressed (*red*) fluctuations are indicated by color, and absolute ΔRMSF values in response of force are indicated by backbone thickness (Fig. 4F). Under pulling, δε exhibits more amplified dynamics than γε, forming a hinge in δ (*red*) and promoting an outward swinging of the semi-detached ε’ (*blue*), further revealing asymmetry between γε and δε.

### Identifying residues critical to *cis*-cooperativity and their distinctive force-responses

To pinpoint the structural determinants of *cis*-cooperativity, we developed a modified occupancy analysis (*i.e.*, ΣOccupancy) that integrated occupancies from CMD and SMD simulations, weighted by the number of connections from that residue to all residues in the binder. Briefly, instead of residue-to-domain occupancy, we quantified residue-to-residue occupancy in a matrix format (fig. S5A). Subsequently, we summed over all possible contacts with the interacting domain (fig. S6, *gray*) to account for one-to-many interactions. This approach broadens the dynamic range of detection and identified 10 and 14% more contacting residues in CMD and SMD, respectively, compared with cryo-EM static structure. We used ΣOccupancy to map high-contact residues at the TCR–CD3 interface, visualized by side-chain color-coding on the structure, highlighting candidate residues with strong interface connectivity (Fig. 4G).

To assess force sensitivity, we calculated the difference between the CMD- and SMD-derived occupancy matrices rather than integrating the CMD and SMD results (fig. S5B). Summing over by row or column yields ΔOccupancy, which in turn identifies force-sensitive residues whose contacts are strengthened (*red*) or weakened (*blue*) by pulling (fig. S6, *red* and *blue*). We mapped these force-sensitive residues on the TCR–CD3 interface by color-coding (Fig. 4H). From a top-down view, force preferentially reduces TCRαβ–CD3γε contacts, whereas TCRαβ–CD3δε contacts are more dynamic. It seems reasonable to suspect that these force-sensitive residues play key roles in mediating force transmission across the TCR–CD3 interface.

### Mutating interfacial residues alters T cell activation and TCR mechanosignaling

Among all *cis*-contact residues, two regions of the TCRβ constant domain (Cβ) stand out by their strikingly opposite mechano-responses. Both the CC’ loop (residues 182-187) and the Helix 4–F strand segment (residues 221-226) is highly conserved and enriched in high ΣOccupancy residues; however, the CC’ loop shows negative ΔOccupancy and the Helix 4–F strand displays positive ΔOccupancy (Fig. 5A). This suggests that force applied from the TCRβ N-terminus to the Cβ domain elicits residue-specific allostery with opposite effects in adjacent structural regions, moving the CC’ loop away from, and the Helix 4–F strand closer to, the CD3s.

**Fig. 5.**
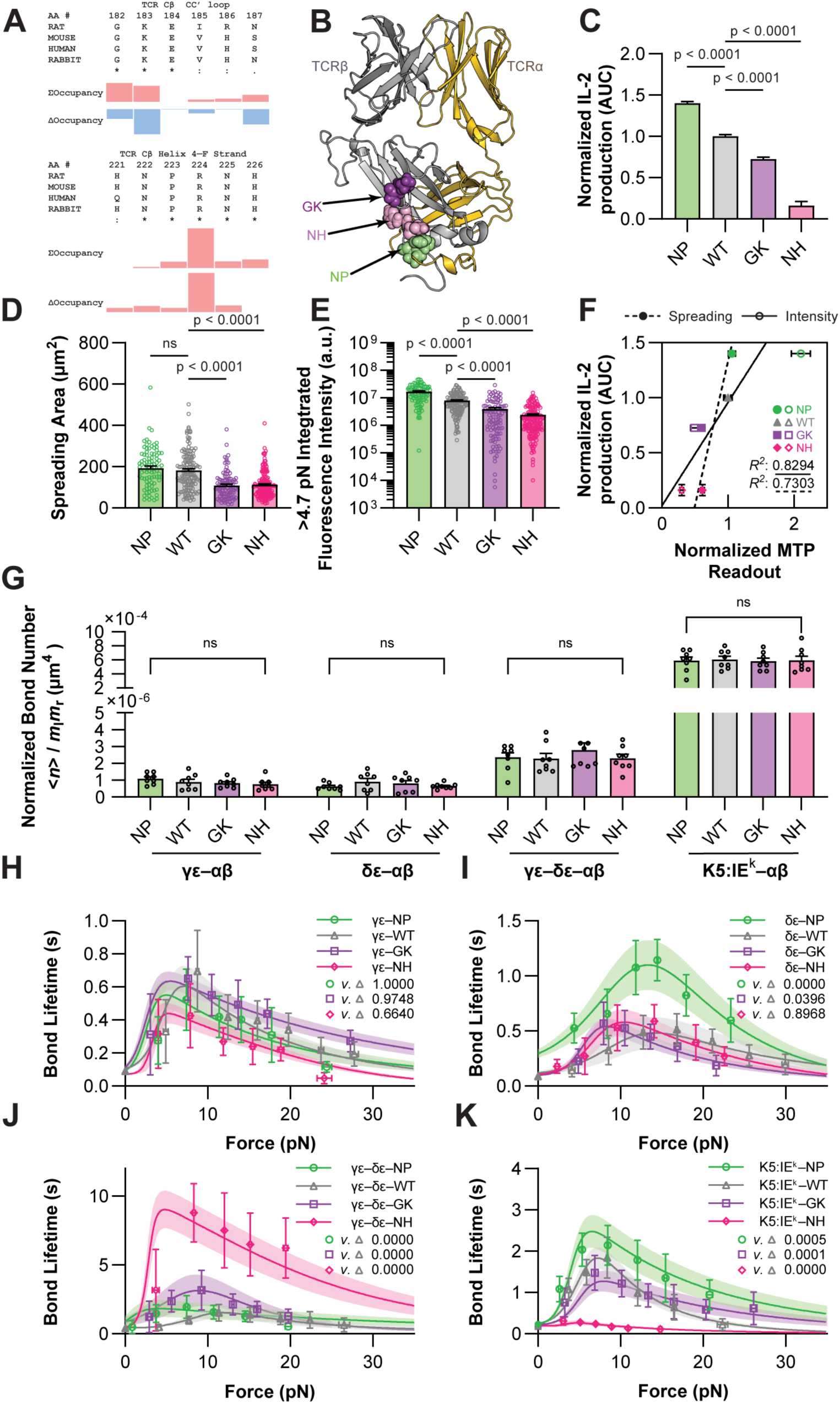
Effects of mutating Cβ contact residues on TCR mechanosignaling, *cis*-interaction affinity and bond lifetime. (**A**) TCRβ local segments with distinct force-responses. Residues in the CC’ loop (*upper table*) and Helix 4 – F strand (*lower table*) of the Cβ domain both contact CD3 (indicated by ΣOccupancy) but respond differently to 175 pN clamping force (indicated by ΔOccupancy). (**B**) TCRαβ structure with double-mutants listed in (A) highlighted on the Cβ CC’ loop and Helix 4-F strand, which contribute to the force-dependent cooperative TCR–CD3 *cis*-interaction according to (A). (**C**) Differential IL-2 cytokine production of cells containing Cβ mutations. The hybridoma expressing 2B4 TCR WT and MTs were stimulated by K5:I-E^k^ displayed on CHO cells. The area under the dose-response curve (AUC), normalized by that of WT, quantifies the relative IL-2 secretion. This data has previously been reported (*26*) and is reformulated here to facilitate experimental design and correlative analysis. (**D** and **E**) Mean ± SEM of spreading areas (D) and integrated fluorescence intensities (E) of individual Jurkat cells (*points*) transduced with WT and MTs TCR (NP in *green*, WT in *gray*, GK in *purple*, NH in *magenta*) when reacting with surface-bound K5:I-E^k^ MTP tethers. The latter signals are proportional to the cumulative number of TCR–pMHC bonds bearing > 4.7 pN cell-generated force to unfold the DNA hairpin tagged to pMHC. Statistical significance levels were assessed by Welch’s t-test assuming non-equivalent SD for indicated pairs. (**F**) Correlations between normalized MTP readouts and IL-2 production. The mean ± SEMs of relative IL-2 AUC in (C) are correlated well with the normalized spreading areas from (D) (*black dashed line*; *closed symbols*) and the normalized integrated fluorescence intensities from (E) (*black solid line*; *open symbols*). (**G**) Normalized average number of bonds of *cis*-interactions between CD3δε (*first group*), CD3γε (*second group*), or CD3δε–CD3γε (*third group*) and TCRαβ WT (*gray*) or NP (*green*), GK (*purple*), NH (*magenta*) MTs. Corresponding ⟨*n*⟩/*m*_l_*m*_r_ of WT and indicated mutant 2B4 TCRs on hybridomas *trans*-interacting with K5:I-E^k^ are plotted for comparison (*fourth group*). Statistics were obtained by comparing any two groups using unpaired Student’s t-tests. ns indicates no significance. (**H** to **J**) Force-dependent bond lifetimes (*points*, mean ± SEM of > 30 lifetime events per force bin; *curves with shades*, mean ± SD of globally fitted bond profiles) of *cis*-interactions of CD3γε (H), CD3δε (I), or CD3δε–CD3γε (J) with TCRαβ WT (*gray*) or NP (*green*), GK (*purple*), NH (*magenta*) MTs. (**K**) Corresponding *τ*(*f*) of WT and indicated mutant 2B4 TCRs on hybridomas *trans*-interacting with K5:I-E^k^ are shown for comparison. This data has previously been reported (*42*) and is replotted here to facilitate correlative analysis and mechanistic understanding. Pairwise significances for mutational effects in (H) to (K) were assessed using chi-square likelihood ratio tests.

We targeted the Cβ CC’ loop and Helix 4–F strand regions, highlighted on the TCRαβ structure (Fig. 5B), by introducing double-alanine mutations, G182A/K183A (GK), N222A/P223A (NP), and N225A/H226A (NH), to alter residues with both strong (large ΣOccupancy) and force-sensitive (high absolute ΔOccupancy) contacts. As previously reported (*26*), mutant mouse 2B4 TCRαβ chains assembled normally with CD3 chains in transduced hybridoma cells, yet altered T cell activation, measured by interleukin-2 (IL-2) secretion upon co-culture with K5:I-E^k^ expressing CHO cells (Fig. 5C, reformatted IL-2 dose-dependent response curves from Ref. (*26*)). Among these, NP acted as a gain-of-function (GOF) mutation, whereas GK and NH behaved as loss-of-function (LOF) mutations.

As a late functional readout, IL-2 production depends not only on TCR–pMHC binding but also on other stimulatory and inhibitory signals along T cell activation pathways. To investigate whether and, if so, how our mutations may affect early TCR mechanosignaling, we measured cell spreading and pulling of Jurkat cells transduced with WT or Cβ mutant 2B4 TCRs on K5:I-E^k^ tagged molecular tension probe 10 min post seeding (MTP; fig. S7A). The MTP assay employs a force-sensitive DNA hairpin that unfolds when tension exceeds a threshold (4.7 pN), yielding a fluorescent signal that reports endogenous force transmitted through the TCR–pMHC bond. As such, the fluorescence signals integrated over time is proportional to the cumulative number of TCR–pMHC bonds per cell. Quantification of the images exemplified in fig. S7B revealed that, T cells with distinctive TCR–CD3 *cis*-constructs spread and pull differently in *trans* through pMHC-specific antigen recognition (fig. S7, C and D). Published studies have demonstrated that these cell-generated forces depend on antigen-induced TCR signaling to activate intracellular force generation and transmission machinery (*50, 74*), therefore can serve as a readout for TCR mechanosignaling. When compared to WT cells, the GOF mutant NP exhibited similar spreading area (Fig. 5D) but significantly increased number of bonds experiencing above threshold forces (110.4 ± 15.7% increase; Fig. 5E). By comparison, LOF mutants GK and NH both showed significantly reduced spreading areas (40.0 ± 5.5 and 38.4 ± 5.0% decreases; Fig. 5D) and integrated fluorescence intensities (50.5 ± 8.7 and 69.9 ± 6.8% decreases, respectively; Fig. 5E). The MTP results correlated strongly with IL-2 data across the variant set (*R*² = 0.73 for spreading and *R*² = 0.83 for pulling, respectively; Fig. 5F), revealing consistent mutational effects in early and late signaling. Jointly, these signaling data underscore the functional importance of TCRαβ–CD3 *cis*-interaction in effectively relaying the TCR–pMHC *trans*-interaction signals for downstream T cell activation.

### Mutating interfacial residues alters catch bond but not affinity of TCRαβ–CD3 interactions

To scrutinize how *cis*-interactions regulate signaling propagation, we measured *cis*-interactions between WT and mutant (MT) N-terminally biotinylated 2B4 TCRαβ with C-terminally biotinylated CD3γε, CD3δε, or both. To ensure the 1:1 ratio of CD3γε and CD3δε constructs, we engineered flexible amino acid linkers to covalently tether the two CD3 heterodimers, thereby avoiding heterogeneous CD3 mixtures (*i.e.*, *in vitro* CD3γε–CD3δε, see Methods). Comparing MTs to WT TCRαβ, statistically indistinguishable normalized numbers of bonds were observed for bimolecular interactions with CD3γε (7.52 – 10.94 × 10^−7^ μm^4^; Fig. 5G, *first group*) and CD3δε (6.25 – 9.11 × 10^−7^ μm^4^; Fig. 5G, *second group*) and much higher but similar-valued were observed for trimolecular interactions with both CD3s (22.75 – 27.82 × 10^−7^ μm^4^; Fig. 5G, *third group*). The cooperative analysis also revealed similar positive cooperativity measures in WT and MT trimers (fig. S8A). Furthermore, *trans*-interactions between complete WT and MT 2B4 TCR–CD3 complexes expressed on hybridoma cells with C-terminally biotinylated K5:I-E^k^ pMHC reported orders of magnitude higher than *cis*-counterparts but near-identical values among each other (5.59 – 5.87 × 10^−4^ μm^4^; Fig. 5G, *fourth group*). These results rule out zero-force kinetics of TCRαβ–CD3 *cis*-interaction and TCR–pMHC *trans*-interaction as a possible reason for the altered T cell signaling by the mutations.

However, under force, more pronounced kinetic differences emerged in terms of the catch-slip bond. Bond lifetime *vs.* force curves for WT and MT TCRαβs with CD3γε were similar (Fig. 5H); so were those with CD3δε, except for the NP mutation (Fig. 5I), which exhibited a markedly enhanced catch bond. Interestingly, when comparing against each other, their force-dependent bond lifetimes demonstrated asymmetry again, as measured by the optimal force where bond lifetime peaks and the catch bond intensities (a metric summarizing force-dependent bond lifetime changes (*42*)), both showing significantly higher values for the TCRαβ mutants interacting with CD3δε than CD3γε (fig. S8, B and C). This validates the results from structural analysis and MD simulations (Fig. 4, C to F), confirming that CD3δε is more force-responsive than CD3γε.

In the case of co-presenting both CD3s, bond lifetime profiles revealed marked mutational effects (Fig. 5J). Notably, the NH LOF TCRαβ mutant formed exceptionally stable bonds with CD3γε–CD3δε, sometimes exceeding 100 seconds, resulting in an average bond lifetime several-fold longer than the WT TCRαβ. Similarly, the GK LOF mutant exhibited an enhanced catch bond with longer lifetimes across the entire force range, whereas the NP GOF mutant displayed a bond profile resembling WT. Recalling our bond measurements for corresponding *trans*-interactions (Fig. 5K, replotted from Ref. (*42*)), the mutational effects on TCR–pMHC *trans*-interactions and TCRαβ–CD3 ECD *cis*-interaction appear to exhibit a reverse correlation: the longer the *cis*-bond, the shorter the *trans*-bond and vice versa.

### Correlating T cell signaling with respective *cis*- and *trans*-interactions of TCRαβ with CD3s and pMHC

We next inspected possible correlations between IL-2 secretion (Fig. 5C) and the kinetic measures of the ECD *cis*-interactions of WT and MT TCRαβs with either CD3γε, CD3δε, or both in the absence and presence of force. For bimolecular *cis*-interactions, as expected from their statistically indistinguishable values across each group, the effective 2D affinities (Fig. 5G, *first two groups*) showed poor predictive power for normalized IL-2 AUC, with steep regression slopes and/or low *R*^2^ values (Fig. 6A). Here, the inverse of the slope indicates the sensitivity-of-prediction, and *R*^2^ reflects the goodness-of-fit (see Methods). Poor predictive powers for T cell function were similarly observed from lifetimes measured at zero force and optimal force (Fig 5, H and I) for both TCRαβ–CD3γε and TCRαβ–CD3δε bimolecular *cis*-bonds (Fig. 6B, *orange and blue*, respectively; fig. S8D). Of note, slightly better correlation with T cell signaling was observed for TCRαβ bonds with CD3δε than CD3γε at both types of lifetime measures, with higher *R*^2^ values and more gradual slopes (Fig. 6B, *dotted vs. solid lines*), potentially because of the differential force-responsiveness.

**Fig. 6.**
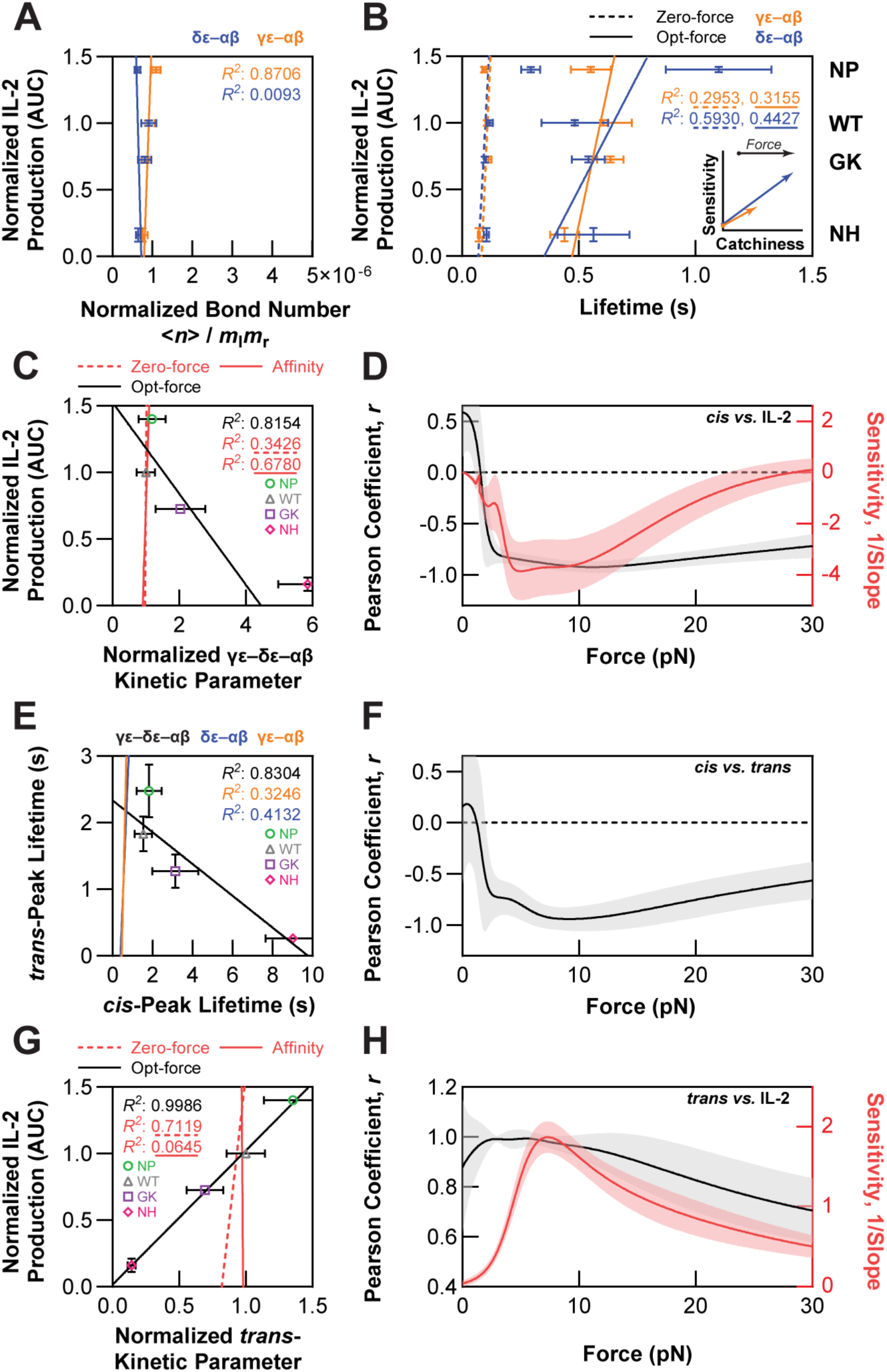
Correlating ECD *cis*-interaction parameters of TCRαβ with CD3γε, CD3δε, or both with T cell function and *trans*-interaction parameters between TCR and pMHC. (**A**) Lack of correlations and/or sensitivities between the IL-2 secretion (Fig. 5C) and effective 2D affinity for αβ–γε (*orange*) or αβ–δε (*blue*) *cis*-interaction (in Fig. 4C, *first* and *second group*). (**B**) Lack of correlations and/or sensitivities between IL-2 secretion and zero-force (*dotted lines*) *vs.* optimal-force (*solid lines*) bond lifetime for αβ–γε (*orange*) or αβ–δε (*blue*) *cis*-interaction (Fig. 5, H and I). *Inset*: Different catchiness and sensitivity of CD3γε (*orange*) *vs.* CD3δε (*blue*). The arrows display the effects of force. (**C**) Inverse correlations between IL-2 secretion and normalized γε–αβ–δε *cis*-kinetics in the presence (but not absence) of force. The mean ± SEMs of relative IL-2 AUC (Fig. 5C) are plotted against the peak lifetimes of ECD *cis*-interactions of WT and MT TCRαβs (Fig. 5J) when both CD3γε and CD3δε are co-presented (*black*). The poor correlations and sensitivities between IL-2 production and zero-force bond lifetime (*red dashed line*) or normalized number of bonds (*red solid line*) for γε–αβ–δε *cis*-interactions are also plotted for comparison. (**D**) Goodness-of-fit and sensitivity-of-prediction of force-dependent bond lifetimes of γε–δε *cis*-interacting with WT and MT TCRαβs in predicting downstream IL-2 responses. The Pearson correlation coefficients (*black*) and inverse of the linear-regression slopes (*red*) between normalized IL-2 AUC and γε–δε–αβ *cis*-catch bonds are plotted for a range of forces (*curves with shades*, mean ± SD). (**E**) Inverse correlations between lifetimes *trans*-bonds and trimeric γε–αβ– δε (but not dimeric αβ–γε and αβ–δε) and *cis*-bonds. The mean ± SEMs of the peak lifetimes of trimolecular *cis*-interactions (Fig. 5J) are plotted against those of *trans*-interactions from WT and MT 2B4 TCRs expressed on hybridoma cells interacting with K5:I-E^k^ (Fig. 5K). The poor correlations and sensitivities between the peak lifetimes of bimolecular *cis*-interactions (*orange*, αβ–γε and *blue*, αβ–δε) with the peak lifetimes of the WT and MT 2B4 TCRs *trans*-interacting with K5:I-E^k^ are plotted for comparison. (**F**) Goodness-of-fit of force-dependent bond lifetimes of γε–αβ–δε *cis*-interacting with WT and MT TCRs in predicting corresponding TCR–pMHC *trans*-force kinetics. The Pearson correlation coefficients of the *cis*-*vs. trans*-bond lifetime are shown for every single force magnitude (*curves with shades*, mean ± SD). Sensitivity-of-prediction is not applicable for this case. (**G**) Inverse correlation between the IL-2 secretion and normalized TCR– pMHC *trans*-interaction kinetics in the presence (but not absence) of force. The mean ± SEMs of relative IL-2 AUC (Fig. 5C) are plotted against the peak lifetimes of *trans*-interactions from WT and MT 2B4 TCRs (Fig. 5K) expressed on hybridoma cells interacting with K5:I-E^k^ (*black*). The poor correlations and sensitivities of the IL-2 AUC with zero-force bond lifetime (*red dashed line*) or normalized number of bonds (*red solid line*) of the TCR–pMHC *trans*-interactions are plotted for comparison. (**H**) Goodness-of-fit and sensitivity-of-prediction of force-dependent bond lifetimes of K5:I-E^k^ *trans*-interacting with WT and MT TCRs in predicting downstream IL-2 responses. The Pearson correlation coefficients (*black*) and inverse of the linear-regression slopes (*red*) between normalized IL-2 AUC and TCR–pMHC *trans*-catch bonds are plotted for a range of forces (*curves with shades*, mean ± SD).

To examine the contribution of cooperativity of the CD3γε–TCRαβ–CD3δε ECD *cis*-interaction to T cell activation, the normalized bond numbers (Fig. 5G, *third group*) and lifetimes at zero force (Fig. 5K, 0 pN) of the trimeric interaction were analyzed, but again showed a lack of functional correlations due to their comparable values (Fig. 6C, *red*), similar to their bimolecular counterparts. In contrast to the poor functional correlations with the peak lifetime at optimal force of the two bimolecular bonds (Fig. 6B, *solid lines*), a significant reverse correlation was observed between the peak lifetime of the cooperative trimolecular bond (Fig. 5K, *optimal force*) and IL-2 production, with a much higher *R*^2^ value and a much flatter slope (Fig. 6C, *black*).

The sharp contrast between the differential predictive powers at zero and optimal forces prompted us to analyze the force effect more rigorously by plotting *vs.* IL-2 AUC, the goodness-of-fit and sensitivity-of-prediction based on WT and MT *cis*-lifetimes under each force, finding that the negative correlation persisted across nearly the entire force range tested (Fig. 6D). Because of the negative correlations, the sensitivity-of-prediction followed the reverse trends of catch bonds, with biphasic behavior most pronounced at optimal force, where force-dependent lifetimes diverged the most between TCR WT and MTs. The Pearson coefficient rapidly approached −1 upon application of force and remained strongly correlated across 5 to 20 pN. These results underscore a mechanotransduction role of the cooperative trimeric *cis*-catch bond.

To explore the interplay between *trans*- and *cis*-interactions, we correlated the peak lifetimes of *trans-* against *cis-*bonds, finding strong anti-correlations with those of the trimer (*R*^2^ ≈ 0.8, Fig. 6E, *black*), but not dimers (*R*^2^ < 0.5, Fig. 6E, *orange* and *blue*). To confirm the force effect on *cis*- and *trans*-coordination, we plotted the force-dependent fitting of *cis*- *vs. trans*-bond profiles for WT and NP, GK, and NH mutants indicated no correlation in the absence of force (Pearson coefficient, *r* ≈ 0 at zero force), and weak correlations in the presence of force for the bimolecular *cis*-bonds (0.33 < *r* < 0.73 for γε, and −0.03 < *r* < 0.82 for δε; fig. S8E), but strong negative-correlations in the presence of force for the trimolecular *cis*-bond (*r* dropped sharply, reaching −1 at 10 pN before rebounding; Fig. 6F). Such correlation analysis reveals an interesting synergy between *trans*- and *cis*-interactions, suggesting a possible feedback mechanism of the TCR mechanosensory machine: from the constant domains, the downstream mechanotransduction region transmitting back to the variable domains, the antigen recognition region.

Consequently, modified *trans*-interactions led to differential IL-2 production in 2B4 T cells expressing WT and MT TCRs, which correlated with the peak lifetime of TCR–pMHC *trans*-bonds, recapitulating our previous finding (*42*). These changes were most apparent under optimal forces and were not evident in 2D affinity or bond lifetimes in the absence of force (Fig. 6G). Similar regression analyses of IL-2 production versus *trans*-lifetime revealed force-dependent goodness-of-fit and sensitivity-of-prediction, showing a nearly mirror-image pattern to Fig. 6D due to inverse-correlated *cis*- and *trans*-bonding synergy (Fig. 6H). Force amplified the mutational effects, driving the greatest divergences in *cis*-followed by *trans*-bond lifetimes near optimal forces, where both the *R*^2^ and 1/slope peaked.

Similar correlative analyses can be performed between early mechanosignaling (Fig. 5E) and respective *cis*- and *trans*-interaction kinetics of TCRαβ with CD3s and pMHC. Since IL-2 and MTP readouts are well correlated (Fig. 5F), we expect results similar to those for IL-2 production (Fig. 6, C and G). Indeed, we found strong correlations between the number of TCR–pMHC *trans*-bonds supporting > 4.7 pN cell-generated forces and the peak lifetimes of the respective TCRαβ–CD3 and TCR–pMHC bonds but are opposite in trends: negative in *cis* and positive in *trans* (fig. S8, F and G, respectively). Collectively, thses kinetic *vs.* signal correlation analyses reveal an allosteric pathway of TCR mechanosensing, where *cis*-TCR–CD3 and *trans*-TCR–pMHC interactions function as synergistic machinery. Modifications at *cis*-interfaces feedback to antigen recognition, resulting in pre-abolished associations or sustained engagements that gate downstream signaling during T cell activation.

## DISCUSSION

Noncovalent *cis*-interactions between the TCRαβ and CD3 chains are essential because they not only hold the octameric TCR–CD3 complex together stably but also provide part of the mechanotransmission pathway that bridges the > 15 nm spatial gap between ligand engagement at the TCRαβ N-termini and ITAM phosphorylation at the CD3 cytoplasmic tails (Fig. 7, *middle right*). Our prior work suggests the biological relevance of this pathway, as mutations targeting TCRαβ–CD3 ECD *cis*-interaction result in respective GOF and LOF phenotypes (*26*). Building on our prior NMR chemical-shift perturbation experiments (*26*) suggesting the possible existence of such interactions, the present study directly quantified the physical binding kinetics of TCRαβ–CD3γε and TCRαβ–CD3δε bimolecular interactions, as well as the CD3γε–TCRαβ–CD3δε trimolecular interaction, both in the absence and presence of force. We performed these measurements in the same membrane-proximal geometry where these interfaces exist, using MAF and BFP assays to overcome technical difficulties, since these interactions are exceptionally weak and fast. Our results show that these ECD *cis*-interactions are not merely structural accessories but are force-bearing, force-regulated bonds, supporting the importance of this mechanotransmission pathway and further elucidating its inner workings. Although our head-to-tail pseudo-*cis* configuration controls ectodomain orientation and enables direct quantification of these weak interactions undetectable by SPR, it does not fully recapitulate native lateral interactions on the cell surface, including confinement by the transmembrane bundle, membrane composition, and intracellular assembly history. The measured force-dependent kinetic properties should therefore be interpreted as ectodomain contributions to TCR–CD3 coupling. Future approaches capable of directly measuring laterally organized interactions in intact membranes will be important for extending these findings.

**Fig. 7.**
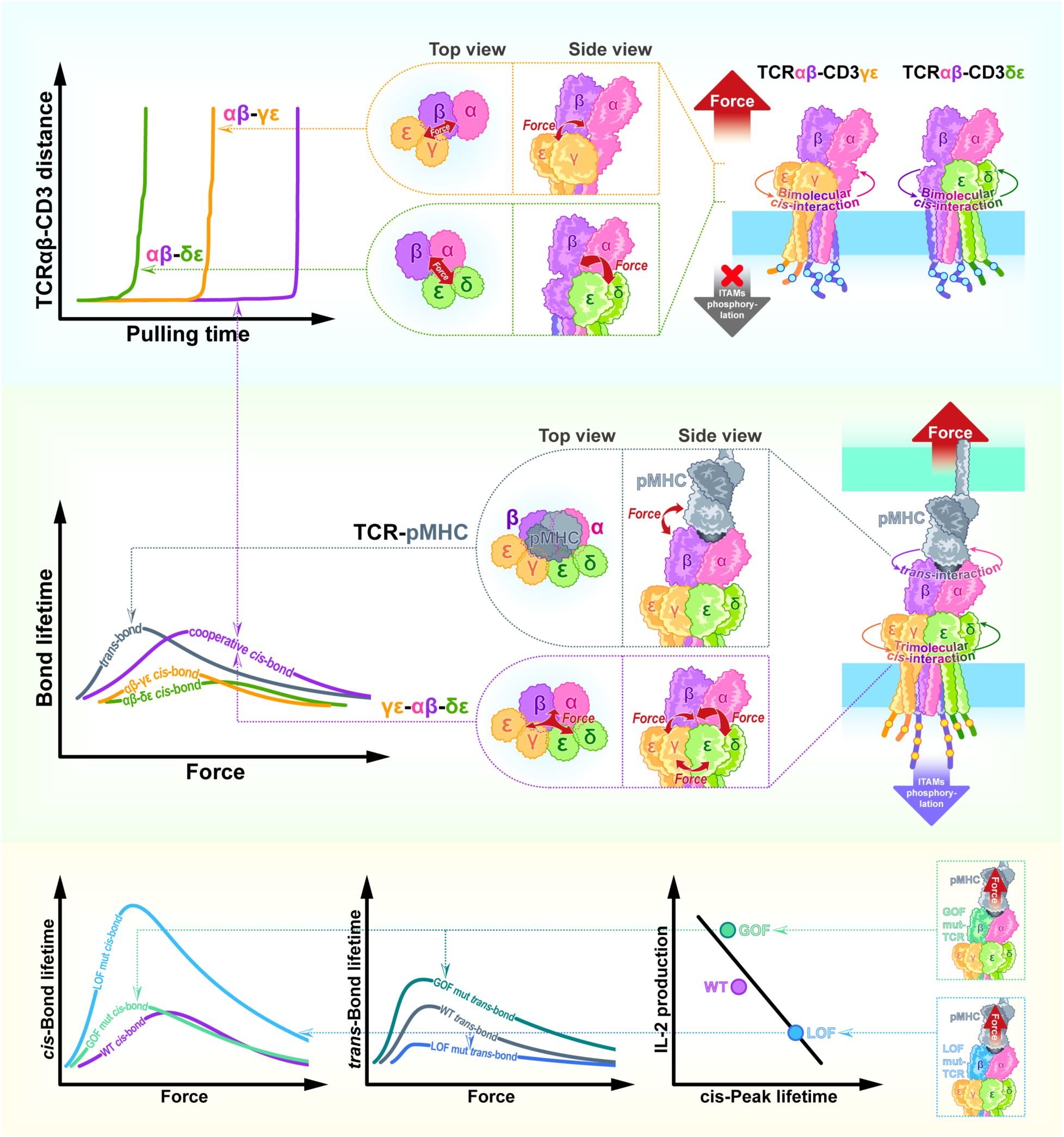
Model for the role of TCRαβ–CD3 ECD *cis*-interaction. *Top row*: Pairwise interactions of TCRαβ with CD3γε and CD3δε (*middle*) exhibit asymmetry, as seen in the SMD simulation (*left*), and are too weak and short-lived to mediate force transmission across the TCRαβ–CD3 interface; hence, they are unable to support TCR triggering (*right*). *Middle row*: While force elicits catch bonds in the pairwise *cis*-interactions (*left*, *orange* and *green curves*), their lifetimes are too short compared to those of the TCR–pMHC *trans*-interaction (*left*, *gray curve*). By comparison, the cooperative trimolecular *cis*-interaction of TCRαβ with both CD3γε and CD3δε (*middle*) generates a greatly enhanced *cis*-catch bond (*left*, *purple curve*) that lasts much longer and matches that of *trans*-interaction of TCR with cognate pMHC. This enables information embedded in the ligand and encoded by force to be transmitted from the pMHC through the TCRαβ to CD3s via interactions of the ECDs, CPs, and transmembrane segments to regulate phosphorylation of the ITAMs in the CD3 cytoplasmic domains (*right*). *Bottom row*: Mutation in the Cβ domain suppresses the *cis*-bond profile (*left*, *light green*) but enhances the *trans*-bond profile (*middle*, *dark green*) and exhibits GOF phenotype (*right*, *green point*), whereas mutation enhances the *cis*-bond profile (*left*, *light blue*) but suppresses the *trans*-bond profile (*middle*, *dark blue*) and displays a LOF phenotype (*right*, *blue point*). This opposite behavior, *i.e.*, a more stable TCR–pMHC *trans*-bond requires a less stable CD3γε–TCRαβ–CD3δε *cis*-bond (to a point yet to be determined), may suggest a mechanotransduction role for ECD *cis*-interactions, which is an interesting topic for future studies.

Key evidence for TCRαβ–CD3 ECD *cis*-interactions to play a mechanotransmission role is the discovery that they form *cis*-catch bonds, indicating that the TCR–CD3 ectodomain junction can function as a mechanically robust and dynamically competent coupling element—capable of transmitting and filtering information under load rather than simply holding the complex together at rest. Importantly, the individual TCRαβ–CD3γε and TCRαβ–CD3δε bimolecular catch bonds are too short-lived to sustain mechanotransmission over the lifetime of an agonist TCR–pMHC interaction (Fig. 7, *top*); yet *cis*-cooperativity provides a solution to this dilemma because strong synergy is generated when CD3γε and CD3δε engage TCRαβ concurrently, they are more likely to form a long-lived trimolecular *cis*-catch bond whose mechanical stability matches that of the TCR–pMHC *trans*-catch bond (Fig. 7, *middle*). While prior assembly studies suggested *cis*-interaction cooperativity among the TM bundles in the TCR–CD3 intra-complex (*17, 75*) since both CD3γε and CD3δε were required for their enhanced assembly with the TCRαβ (*25*), the present work qualitatively demonstrated and quantitatively evaluated the ECD trimolecular binding cooperativity, using a kinetic approach that methodologically follows our previous analysis of positive cooperativity in the TCR–pMHC–CD8 (*50, 76, 77*) and TCR–pMHC–CD4 (*46*) trimolecular interactions, as well as the negative cooperativity between two distinct sets of *trans*-interactions: TCR–pMHC–CD8 and PD-1–PD-ligands on the T cell membrane (*78*). Note that our measurements are likely underestimated because we measured the TCR and CD3 ECDs under pseudo-*cis* configurations without TM bundling, which would place them in the most appropriate orientations, especially when CD3γε and CD3δε were captured randomly as CD3m on the beads. In addition to biophysical measurements, CMD and SMD simulations provide corroborating evidence for strong synergy. These include differences in contact areas, free energies, hydrogen bonds, dissociation times, rupture forces, dynamic footprints, and contact residues between the ECDs of TCRαβ and two CD3s separately or concurrently. Across multiple complementary experimental and computational metrics, our results provide strong evidence for the quantification of positive cooperativity in TCR–CD3 ectodomain *cis*-interactions, as the integrated response of the complex exceeded that predicted from its individual components. Specifically, the whole trimolecular interaction forms more H-bonds with larger footprints and stronger contacts that are energetically more favorable, mechanically more stable, and temporally longer-lasting than the sum of the two bimolecular interactions. Conceptually, cooperativity converts two weak, transient tethers into an integrated “mechanical clutch”: force preferentially stabilizes the coupled state, enabling sustained load sharing and extending the time window for downstream signaling to commit.

The ability of the cooperative TCR–CD3 ECD *cis*-catch bond to support force for times that match the lifetime of cognate TCR–pMHC *trans*-catch bond implies their importance in signal relay (Fig. 7, *middle*). This may be significant considering our current view of the *trans*-catch bond in TCR mechanosensing, which is further supported by the MTP data, suggesting that TCR–pMHC catch bond facilitates TCR triggering, enhances TCR signaling, and amplifies antigen discrimination, providing a greater predictive power for T cell response beyond that of binding affinity in the absence of force (*9, 39, 79*). By comparison, γδTCR–CD3 complexes lack interactions among ECDs with flexible CPs (*80–82*); correspondingly, γδTCR–ligand *trans*-interactions form slip bonds and lack mechanosensing (*12*). This contrast highlights the importance of the cooperative trimolecular *cis*-catch bond revealed here, which must contribute, at least in part, to the “mechanical conduit” with the needed structural connectivity among various αβTCR–CD3 chains important for signaling. Such connectivity must be critical for supporting force, regardless of whether applied externally by pMHC or generated endogenously and exerted through CD3 cytosolic domains. An interrelated dual role may be envisioned for the cooperativity-reinforced *cis*-catch bond: First, force does not merely test bond strength; it selects bond state. Catch behavior implies an underlying force-favored bound conformation (or pathway) that is accessed more frequently or stabilized under loads. Second, because the cooperative *cis*-catch bond is maximally stabilized within a defined narrow force regime, the TCR–CD3 junction can act as a “force band-pass filter,” preferentially transmitting signals within the physiological force window and rejecting either insufficient load (no stabilization) or excessive load (slip-driven rupture). In this view, the mechanistic role of *cis*-cooperativity is not simply to strengthen the complex, but to shape how mechanical information is encoded in bond lifetimes and, thus, signaling probability. Therefore, our findings not only support the TCR mechanotransduction hypothesis (*9, 39*), but are also consistent with the kinetic proofreading model (*83, 84*).

Further support for this view comes from observed fluctuations in the CD3s that are inversely correlated with the cooperatively enhanced contact stability, consistent with dynamic allostery rather than purely elastic linkages within both TCRαβ and CD3 ECDs. NMR (*32, 33*) and CMD (*33, 41*) studies have observed long-range allosteric communication across the TCRαβ ECD upon ligand engagement, manifesting as chemical shift changes and increased dynamics at the CD3 binding site, suggesting regulatory capacity via distant conformational coupling. Dynamics of CD3s have been hypothesized to be the primary drivers of cross-membrane signal transmission, based on their more structured CP regions compared to the flexible TCR anchors (*85*). Although early cryo-EM (*29, 30*) and photo-crosslinking (*27*) studies failed to detect a different stable conformation upon pMHC or activating antibody ligation in the absence of force, a recent cryo-EM study observed distinct conformations of the TCR–CD3 complex preserved in native states using nanodiscs (*35*). The conceptual connection is that the cooperative *cis*-catch bond provides a plausible means to stabilize and mechanically couple such large-scale ECD rearrangements: weak pairwise contacts would be insufficient to sustain coordinated motion, whereas a cooperative *cis*-catch bond could maintain quaternary integrity while allowing force to bias transitions between compact and open ensembles. Thus, our findings are consistent with the allostery hypothesis (*18, 86*).

Also consistent with the view of mechanically regulated dynamic allostery is the finding of asymmetric responses of the two CD3 ECDs, arising from their dissimilar molecular architectures and distinct *cis*-interactions with the TCR α and β chains (Fig. 7, *top*). The Cβ FG loop has been suggested to be important for TCR mechanosensing, partly because of its proximity to the CD3γ ECD, which may restrict domain motions (*40, 87, 88*). Additionally, CD3δε exhibits heavier glycosylation than CD3γε (*57, 89*), suggesting a tighter association of CD3γε with TCRβ than CD3δε. Here, we found that across SMD and BFP (but not CMD and MAF) measurements, CD3δε is more force-responsive and undergoes greater contact reshaping and more vibrant fluctuations, whereas CD3γε maintains a more invariant interface with TCRαβ and appears mechanically more stable. This division of labor suggests a novel conceptual model: CD3γε behaves as a stabilizing strut that preserves the coupled architecture under load, while CD3δε behaves as a compliant hinge that accommodates force-induced rearrangements. Such asymmetric compliance provides a plausible way to meet two competing requirements of mechanotransduction: the complex must remain connected long enough to transmit force-encoded information, yet remain dynamic enough to permit the conformational changes required for signal propagation that ultimately transition to tail exposure and ITAM phosphorylation. Indeed, the ECD asymmetry revealed here may be related to the structural (*35*) and functional (*90*) differences of CD3 chains during TCR triggering. Our findings of the asymmetric behaviors of ε chains suggest an intriguing hypothesis for further investigation: Upon TCR–pMHC binding, release of the cytoplasmic tails of γ- and δ-associated ε’s may also be asymmetric, with the former being kept in a constrained, inactive conformation by binding to the inner leaflet of the plasma membrane, while the latter being exposed to the cytoplasm, ready to be phosphorylated by Lck to initiate TCR signaling.

In this work, we used mutagenesis to perturb the ECD *cis*-bonds within the TCR–CD3 assembly to further elucidate their potential role in mechanotransduction. We examined three double alanine mutations in the TCRαβ constant domains because these residues have high dynamic contacts and force responses with the CD3s, and they show GOF (NP) and LOF (GK and NH) effects as measured by IL-2 production in antigen-stimulated T hybridoma cells (*26*). The MTP assay has provided supporting evidence for the mechanotransduction role of the TCR–CD3 ECD *cis*-bonds by showing that mutations known to perturb TCRαβ–CD3 *cis*-interaction also affect TCR mechanosignaling. Since none of these mutants altered their effective 2D affinities with either CD3 or the normalized average number of bonds with both CD3s when measured without force, the likely biophysical parameter that may impact pMHC-induced TCR signaling must be the force-elicited *trans*-catch *vs.* slip bonds. Indeed, a strong correlation was observed between the number of TCR–pMHC *trans*-bonds transmitting >4.7 pN cell-generated forces and the kinetic properties of these *trans*-bonds in the presence (but not absence) of force (fig. S8G). These results are consistent with the increasingly accepted view that catch and slip bonds are superior predictors of T cell activation, as they link altered *cis*-interface mechanics to early TCR mechanosignaling, as well as to the later integrated IL-2 response. Furthermore, the consistent TCR mechanosignaling results support our mechanosensor model of the TCR–CD3 complex as an intricate yet elegant nanomachine whose various parts and intramolecular interactions contribute to transmitting mechanically encoded information across its structure.

We observed a lack of mutational effect in bimolecular catch bonds except for the NP mutant, which forms a much more pronounced catch bond with CD3δε but not CD3γε. Again, this force-dependent asymmetry may be related to the asymmetric behavior of the two ε cytoplasmic tails, for the more adaptive TCRαβ–CD3δε *cis*-bond may serve as the preferred path of force transmission to enable the larger share of total force through the ε’ chain, facilitating the release of its cytoplasmic tail for signal initiation. Hypothetically, mechanotransmission requires the TCR–CD3 complex to have a mechanically integrated architecture with directional constraints, achieved by asymmetric coupling, to allow different ligands to tip the balance among pre-existing states of an energetically metastable structure, enabling force-regulated conformational changes to propagate allosterically. Thus, the Cβ mutants further sharpen the mechanistic picture by separating equilibrium binding from mechanical coupling. Despite leaving force-free *cis*-affinity/bond number largely unchanged, these substitutions remodel the force-dependent lifetime landscape—most prominently for the cooperative trimolecular *cis*-catch bond—demonstrating that the functional lever resides in force-activated dynamics rather than in static affinity. In other words, these mutations tune the mechanical impedance of the TCR–CD3 junction: they alter how the interface redistributes force and stabilizes (or destabilizes) mechanically favored bound states. In light of the recent strong interest in engineering TCR catch bonds for cancer immunotherapy (*67, 91*), our findings also provide a practical design principle: mechanotransduction can be engineered by targeting residues that modulate the “force-response” of *cis*-coupling, without necessarily perturbing antigen recognition chemistry.

Surprisingly, the GOF mutation suppressed the trimolecular *cis*-catch bond, whereas both LOF mutations enhanced it, showing that the GOF mutant enables, whereas the LOF mutants disable, TCRαβ synergistic associations with CD3γε–CD3δε. This apparent paradox is explained by force-stabilizing and force-destabilizing contacts in our MD simulations, whose removal enhances and restricts CD3 flexibility, respectively, thereby amplifying and attenuating downstream signaling. Indeed, this explanation is consistent with cryo-EM and crosslinking studies, which show that TCR mutants that strengthen the TCR–CD3 interface paradoxically impair T-cell function (*27*), underscoring the functional relevance of CD3 motions.

Previously, we found the same mutations produced a positive correlation between T-cell function and *trans*-bond profile: the GOF mutant TCR (NP) formed a stronger catch bond than WT, whereas the LOF mutants (GK and NH) formed a weaker catch bond or even a slip bond, supporting the predictions from our structural and biophysical model of TCR–pMHC bond profile (*42*). The model suggests that domain stretching, tilting, and rotating near the hinges of the Vα/Vβ and Cα/Cβ domains facilitate *trans*-catch bond formation. A striking finding is the inverse relationship between the cooperative *cis*-catch bond and TCR signaling, along with the anti-correlation between the *cis-* and *trans*-bond profiles (Fig. 7, *bottom*). These trends suggest a tradeoff between internal complex stability and external ligand sensitivity: excessive stabilization of the intra-complex *cis*-junction may “over-clutch” the receptor, restricting the structural compliance that supports optimal *trans*-catch bond formation and antigen discrimination, whereas a more labile *cis*-junction may permit the conformational freedom needed to maximize *trans* mechanosensing— up to the point where connectivity becomes limiting. Alternatively, the dynamic turnover of TCR–CD3 assemblies at the immunological synapse requires cycles of engagement and disengagement; longer-lived *cis*-bonds may slow this turnover, limiting the rhythmic signaling pulses necessary for sustained T cell activation—a mechanism that is non-exclusive with the conformational compliance model described above (*92*). Thus, the unexpected results from this work actually validate our previous model: the weaker the *cis*-catch bond, the more flexible the TCR–CD3 complex, the stronger the *trans*-catch bond, and the greater TCR signaling and T cell function. This view is also consistent with previous studies, finding that LOF and GOF mutations in TM residues correspond to those that stabilize and destabilize the TCR–CD3 complex, respectively (*18, 93*). We therefore propose that TCR triggering requires not maximal *cis*-stability per se, but a balance between connectivity and mobility, such that force can be transmitted while motion remains permissible.

## MATERIALS AND METHODS

### Proteins, cells, and antibodies

A biotinylation sequence (GLNDIFEAQKIEWHE) was added to the N-terminus of ECD constructs of 2B4 TCRαβ and the C-terminus of human CD3γε, CD3δε, and CD3γε–CD3δε subunits by PCR. For the single heterodimer CD3γε and CD3δε, a 26-residue peptide linker, C-GSADDAKKDAAKKDDAKKDDAKKDGS-N, was engineered to connect different CD3 chains (*94*). For the CD3γε–CD3δε heterodimer–heterodimer, a different 25-residue peptide linker, C-GSSPNSASHSGSAPQTSSAPGSQGS-N, was added to join the two dimers (*95*). Proteins were expressed in One Shot^TM^ *E. coli* BL21 (DE3) (Invitrogen) as insoluble inclusion bodies, refolded, and purified as previously described (*26*). Inclusion bodies for I-E^k^ α (with C-terminal biotinylation sequence) and β chains were produced in One Shot^TM^ *E. coli* BL21 (DE3), refolded with K5 peptide (ANERADLIAYFKAATKF), and purified as described previously (*96*). For all proteins, purified fractions were biotinylated using a BirA protein ligase kit (Avidity LLC). The biotinylated proteins were further purified by gel filtration (S200, GE Life Sciences) in PBS (pH 7.4). The proteins were tested for biotinylation efficiency by gel-shift analysis using SDS-PAGE electrophoresis after incubation with excess streptavidin.

Mouse 58^−/-^ T cell hybridoma cells (*97*) expressing mouse CD3 but not TCRαβ were a generous gift from Dr. Bernard Malissen (Centre d’immunologie de Marseille-Luminy, France). 58^−/-^ cells were transfected with mutant mouse 2B4 TCR constructs through retroviral transduction and cultured as described previously (*26*). The transduced cells were stained with phycoerythrin (PE)-conjugated anti-mouse CD3ε (clone 145-2C11, eBioscience) and allophycocyanin (APC)-conjugated anti-TCRβ (clone H57-597, eBioscience) mAbs, then sorted for dual expression of CD3 and TCR. TCR-deficient Jurkat cells (J.RT3-T3.5, ATCC) were transfected with mutant 2B4 TCR through retroviral transduction and cultured as described previously (*27*). The transduced cells were stained with PE-conjugated anti-human CD3ε (clone OKT3, eBioscience) and pacific blue (PacBlue)-conjugated anti-TCRβ (clone H57-597, BioLegend) mAbs, then sorted for dual expression of CD3 and TCR. The sorted cells were expanded for 6 days and quantified for TCR and CD3ε expression.

### Flow cytometry and site density

Samples were stained using 10 µg/ml (as indicated below) of antibodies in 100 µl of FACS buffer (PBS or Hanks Ca^2+^/Mg^2+^ free, 2% FBS, 5 mM EDTA, and 0.1% NaN3) for 30 min at 4 °C. Subsequently, samples were washed twice in 500 µl and then resuspended in 300 µl of FACS buffer. High-performance analysis and sorting were conducted using BD FACSAria™ III, and then flow cytometric data were processed using FlowJo™ Software. PE-conjugated anti-mTCRβ (clone H57-597, BD Biosciences), anti-mI-E^k^ (clone 14-4-4S, BD Biosciences), and PE-Cy7-conjugated anti-mCD3ε (clone 145-2C11, BD Biosciences) were used for *trans*-interactions. PE-conjugated anti-mTCR Vα11.1, 11.2 (clone RR8-1, BioLegend), anti-hCD3ε (clone UCHT1, BioLegend), isotype rat IgG2b,κ (clone RTK4530, BioLegend), isotype mouse IgG1,κ (clone MOPC-21, BioLegend) were used for *cis*-interactions.

### Micropipette adhesion frequency assay

The MAF assay for measuring 2D affinity has been described previously (*59, 60*). Briefly, human RBCs were isolated from whole blood obtained from healthy donors using a protocol approved by the Institutional Review Board of the Georgia Institute of Technology. RBCs were biotinylated with various concentrations of biotin-X-NHS linker (Millipore/Sigma) to achieve different levels of surface functionalization. Then, the biotinylated RBCs were incubated with a saturating amount of tetrameric streptavidin (Thermo Fisher) and the biotinylated protein of interest for 30 min at room temperature. Based on available biotin sites, the protein was fully coupled to RBCs via biotin-streptavidin-biotin conjugations, and the resulting surface density was quantified by flow cytometry.

An experiment chamber was assembled by sandwiching coverslips onto short-edge metal spacers, leaving long-edge gaps open for inserting solutions and reagents. The channel bridging two metal spacers was created by filling the center space with Leibovitz’s L-15 Medium (Gibco) and sealing the solution with mineral oil. The chamber was mounted on the stage of an inverted microscope (a 40×/NA 0.75 objective with a 4× TV tube, Nikon, Eclipse Ti) and imaged at 200 frames per second (Basler, Ace 2). RBCs, beads, and hybridoma cells were loaded into different regions of the channel; the forged micropipettes (Sutter Instrument, P-97) were inserted from each side of the openings to form opposing alignments.

Using suction pressures from the manometer systems, a pair of RBCs (or an RBC and a bead or a hybridoma cell), bearing receptor and ligand, respectively, were captured by two opposing capillary micropipettes (Fig. 1A). The piezo-driven target RBC on the right was brought into contact with the stationary probe RBC on the left in the repeated “approach-contact-retract” testing cycles. Binding *vs.* non-binding events were sampled at a constant contact time and contact area: upon retraction, RBC elongations indicate binding as 1, and no deformations indicate non-binding as 0. The adhesion frequency (*P*_a_ = ⟨*s*⟩) is the mean of the obtained Bernoulli sequence (*i.e.*, {*s*_1_, *s*_2_, *s*_3_, ⋯, *s_n_*}), in other words, the probability that binding events occur.

Due to the ultra-low affinities, the *cis-*bond formations are *i.i.d.* and rare. Thus, the probability of forming *n* bonds follows the Poisson distribution, *p_n_* = (⟨*n*⟩*^n^*/*n*!) exp(−⟨*n*⟩), with a single parameter, ⟨*n*⟩, the mean number of bonds formed during contact time. As a result, the adhesion frequency is expressed as,

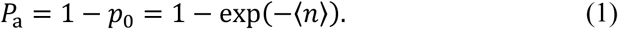

To further relate the characteristic ⟨*n*⟩ to pairwise *cis*-binding kinetics, we established the kinetic differential equation that is governed by the second-order forward and first-order reverse reactions,

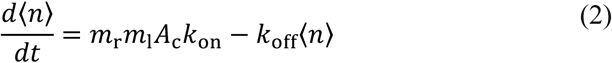

where *m*_r_ and *m*_l_ (μm^−2^) are site densities of receptors and ligands on the RBC surfaces, *A*_c_ (μm^2^) and *t* (s) are contact area and time; *k*_on_ (μm^2^ s^−1^) and *k*_off_ (s^−1^) are 2D on-rate and off-rate, whose ratio gives 2D affinity *K*_a_ = *k*_on_/*k*_off_ (μm^2^). Clearly, one of the solutions to Eq. 2 is,

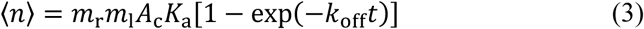

whose integration constant is zero, assuming there is no bond initially. Because of the rapid off-rates of *cis*-interactions, the exponential term vanishes at steady-state with a long contact time (2 s). As a result, effective 2D affinity *A*_c_*K*_a_ can be simply derived from *P*_a_ by,

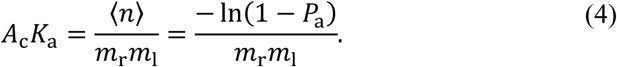

Here, *m*_r_ and *m*_l_ were determined by calibrating the mean fluorescence intensity (MFI) of a protein of interest fully stained with a PE-tagged antibody against the MFI of PE quantitation beads (BD Biosciences) by flow cytometry.

### Biomembrane force probe assays

The BFP thermal fluctuation (*60, 62*) and force-clamp (*43, 50*) assays for measuring bond lifetime in the absence and presence of force, respectively, have been described in detail previously. Briefly, isolated human RBCs were treated and functionalized to form ultra-sensitive mechanotransducers. Probe RBCs were fully biotinylated with a saturating amount of biotin-PEG-NHS (JenKem), then inflated into hypotonic states by nystatin (Thermo Fisher). Next, the probe beads were prepared for dual roles: to attach to the RBCs and to display the protein of interest. In short, streptavidin-maleimide (Sigma) was clicked onto thiol-functionalized glass beads (MPTMS, Thermo Fisher) overnight at 4°C. Afterward, the biotinylated protein-of-interest was partially coupled to the streptavidin beads (90 minutes at room temperature), leaving enough vacancies for RBC attachments through biotin-streptavidin interactions. Lastly, either beads, RBCs, or cells were used as the targets, on which the binding partners were either chemically conjugated or naturally assembled for the probe to test against.

A similar experiment chamber (see MAF section) was prepared prior to loading the probe RBCs, probe beads, and targets. In addition to the two opposing micropipettes, a third helper pipette was inserted to assist probe assembly: capturing smaller probe beads and attaching them to the apex of RBCs. The microscope was set up the same way, but in addition to Basler visualization, another high-speed CCD camera (Mako G040B, Allied Vision) was used to track deformations in the probe RBCs. To do so, a 60-line strip window across the probe was imaged at 1,200 frames per second, where a homemade dual-edge tracking program (LabVIEW, Texas Instruments) scanned the real-time bead positions relative to the reference on the pipette (*i.e.*, the axial length of probe RBCs). This drift-prone design enables ultra-stable and ultra-sensitive micro-manipulations with 2 nm spatial and 1 ms temporal resolution.

#### 1. Thermal fluctuation assay

This assay is based on the equipartition theorem (*62, 65*), where the RBCs elastic energy is quadratic under small deformations,

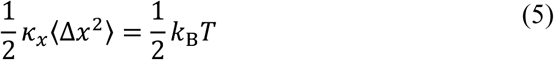

where *κ_x_* is the spring constant of RBC’s simple harmonic motion, and *k*_B_ is the Boltzmann constant that scales absolute temperature to energy. The union of the target with the probe, via bond formation, dampens its Brownian motion, equivalent to adding a parallel spring that stiffens the system with a ramped-up spring constant. Whereas bond breakage leads to the restoration of *κ_x_*. Because of the anti-correlation between *κ_x_* and (Δ*x*^2^⟩, we pinpointed bond association and dissociation events by sudden decreases and increases in the variance of probe displacements, respectively.

During measurements, we adjusted the manometer system to lightly grab the probe RBC with the left micropipette, keeping the RBC ultra-soft with small *κ_x_* to enable sensitive detection. Subsequently, the probe bead was aligned and pushed against the apex of probe RBC, where biotin-streptavidin firmly fused them to assemble the complete mechano-sensing probe (Fig. 1B, *left*). Then, we brought the target into proximity with the probe, leaving a tiny gap between the target’s receptor-presenting and the probe bead’s ligand-presenting surfaces. Hence, the thermally driven Δ*x* occasionally closed the gap to allow ligand-receptor binding. If bonds occurred, the durations between the onsets of (Δ*x*^2^⟩ reductions and the moments of (Δ*x*^2^⟩ resumptions (*i.e.*, sequence of events from the exponential random variable, {*t*_1_, *t*_2_, *t*_3_, ⋯, *t_n_*}) were sampled, whose mean reports the ligand-receptor lifetime, *τ* = ⟨*t*⟩. Note that the adhesion frequency was kept low enough to ensure limited second bond formation before breaking the first bond.

To calculate the off-rate, *k*_off_ of pairwise interactions from sampled lifetime events, the kinetic differential equation of first-order reverse reaction is simplified by setting *k*_on_ = 0 (Eq. 2) as,

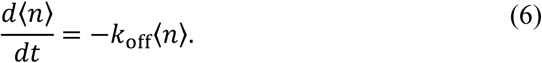

Its solution is scalable with the initial number of bonds, ⟨*n*_0_⟩, and reexpressed as the time-dependent survival probability, *P*_s_(*t*),

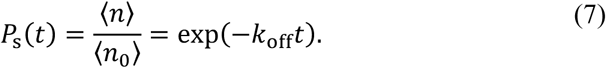

As a result, *P*_s_(*t*), the number of events lasting longer than *t* normalized by total lifetime events, starts from point (0, 1) and decays exponentially. After rearranging lifetime events chronologically, the linear regression slope of survival probability *vs.* time on a logarithmic scale estimates *k*_off_.

#### 2. Force-clamp assay

The force-clamp assay utilized the same BFP micropipette apparatus, probe setups, and coating strategies as the thermal fluctuation assay (Fig. 1B). Nonetheless, instead of passively monitoring lifetime events without interference, we precisely quantified and interactively manipulated the force levels exerted on the ligand-receptor bonds to obtain the force-dependent lifetime, *τ*(*f*). The probe, as a harmonic oscillator, obeys Hooke’s law,

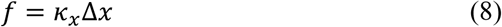

where *κ_x_* converts the axial displacements into the force applied by the spring. To fine-tune probe stiffness, the suction pressure (Δ*p*) on probe RBCs modulates the hypotonic RBC membrane tension, which, in turn, relates to the spring constant by a geometric scaling factor as,

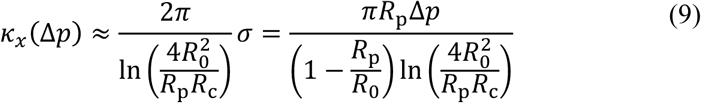

where *σ* is the membrane tension, and the characteristic measures of the probe, *R*_0_, *R*_p_, and *R*_c_, are radii of the outer RBC sphere, the inner RBC cap, and the bead-RBC circular contact, respectively. Typically, the designated Δ*p* sets *κ_x_* ≈ 0.3 pN/nm, which establishes the ultra-sensitive force sensor at a resolution of ± 5 nm in *x*-deformations.

During experiments, once fine-tuned to the specified stiffness, we utilized a high-precision piezoelectric linear actuator (Physik Instrumente) to micro-manipulate the target, allowing it to undergo “approach-contact-retract-hold-return” testing cycles against the stationary probe. Based on the instantaneous *f*(*t*), automated subprograms (LabVIEW, Texas Instruments) made interactive decisions about the target’s sub-nanometer movements. The target **approached** the probe till *f*(*t*) dropped below zero. Next, the target was stopped at *f*(*t*) ≈ −25 pN and **contacted** the probe to facilitate ligand-receptor associations for a fixed period of 0.5 s. At its end, the piezo **retracted** the target from the probe with two outcomes: a non-binding event was indicated by (*f*(*t*)⟩ ≈ 0, and a binding event was detected by (*f*(*t*)⟩ ≫ 0 at the moment of separation (Fig. 1F). If the ligand-receptor formed a bond and the ramping-up force reached an expected level, *f*, the target was **held** in place to maintain the constant Δ*x*, such that the tensile force on the bond was clamped until the spontaneous dissociation occurred (*i.e.*, the sudden decline of *f*(*t*)). Later, the piezo **returned** the target to its origin to reinitiate the next testing cycle. Among 100s of cycles tested on a pair of targets and probes, lifetime events (*i.e.*, 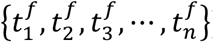) were sampled as the time intervals between the instants of *f*(*t*) ramped up to the desired magnitudes, and the moments of *f*(*t*) suddenly dropped to zero. The mean lifetime reports the force-dependent ligand-receptor lifetime, *τ^f^* = (*t^f^*), where the superscript denotes force clamping of *f*. To ensure that most interactions represented single-bond events, we adjusted the site densities of the molecules coated on the beads to keep the adhesion frequency below 20%.

### Cooperativity analyses

We used multiple metrics to assess the cooperativity of *cis*-interaction, such as 2D affinity and force-dependent bond lifetime. When CD3γε and CD3δε were co-presented, comparisons between the actual measurements and expected values under independence revealed cooperativity of the TCR–CD3 mixture. To formulate a criterion and a quantitative index for cooperativity, we derived the general form as,

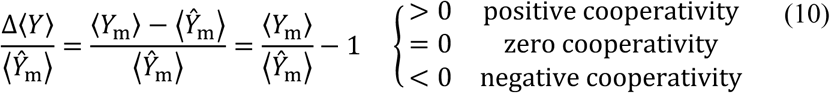

where *Y* is an arbitrary random variable about *cis*-interaction, and *Y*_m_ and *Ŷ*_m_ are the obtained and expected values from the CD3 mixture, respectively. Zero cooperativity indicates independent binding, while positive or negative cooperativity indicates dependent binding, with the magnitude of cooperativity quantifying the degree of dependence. We used three types of cooperativity analyses to evaluate *cis*-binding between TCRαβ and CD3γε and CD3δε mixtures using *in vitro* MAF, BFP, and *in silico* SMD dissociation assays. Under the assumption of independent bindings, the expected values for corresponding assays are listed below,

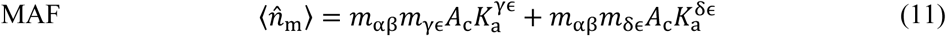

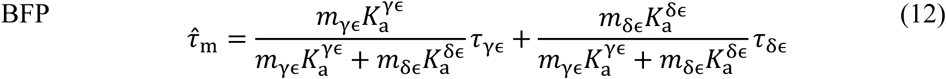

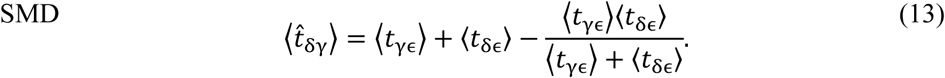

Here, *K*_a_, *τ*, and ⟨*t*⟩ in expressions are characteristic kinetic parameters of pairwise *cis*-interactions quantified from corresponding TCRαβ–CD3γε and TCRαβ–CD3δε assays, while *A*_c_ and *m* are controllable factors. Note that *τ* and ⟨*t*⟩ distinguish lifetime quantification using different approaches: the former population study reports the ensemble average of multiple association states (subscript m), and the latter computer study is based on the single trimeric state (subscript δγ).

### Derivations of the expected value of trimeric dissociation lifetime

The expected values for the first two cooperativity analyses are straightforward, and we only show the derivations for Eq. 13 in detail here. The third cooperativity analysis is based on dissociation waiting times during SMD simulations. Again, this differs from the population means of BFP lifetime measurements because we simulated a single-molecular construct at a time, either TCRαβ–CD3γε or TCRαβ–CD3δε, or CD3γε–TCRαβ–CD3δε complexes. Recall that the survival probabilities for the pairwise associations of TCRαβ–CD3γε and TCRαβ–CD3δε are,

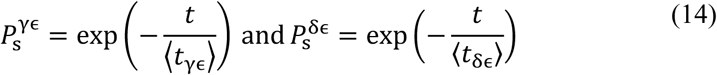

respectively, where (*t*_y**ε**_) and (*t*_δ**ε**_⟩ are their corresponding average waiting times until dissociation. Assuming the CD3γε and CD3δε dissociations from the concurrently bound states are independent, and the presence of high loads prevents any rebinding upon dissociation, the cumulative joint probability for TCRαβ to dissociate from both concurrently during (0, *t*) is,

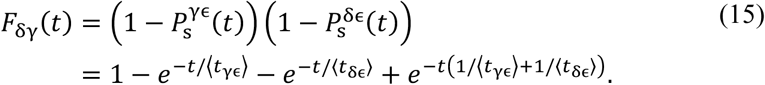

The corresponding probability density is,

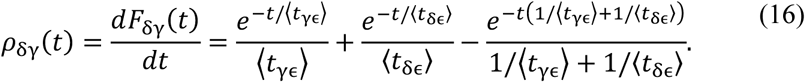

Thus, the expected value of trimeric dissociation lifetime under the non-coop assumption is,

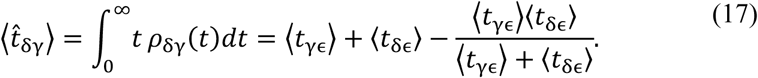

### Kinetic modeling of dual-species interaction

To model the dual-species scenario, in addition to the pairwise interactions in the single-species scenario, only the CD3γε–TCRαβ–CD3δε trimolecular interaction is allowed as a concurrent binding state. As such, for simplicity, the total TCRαβ bonds with CD3m include only three subpopulations.

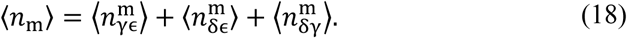

where *n* with different subscripts denotes the number of bonds for the TCRαβ–CD3γε, TCRαβ–CD3δε bimolecular interactions, and the CD3γε–TCRαβ–CD3δε trimolecular interaction, respectively. Here, the concurrent mode exhibits unique kinetics, redistributing the probabilities of dimeric and trimeric bond formation.

The competitive kinetics between the pairwise and concurrent TCRαβ–CD3s *cis*-interactions follow the two-step model. In the first step, the TCRαβ interacts with either CD3γε (kinetic rates *k*_1_ and *k*_–1_) or CD3δε (kinetic rates *k*_2_ and *k*_–2_) to form TCRαβ–CD3γε or TCRαβ–CD3δε bimolecular bonds. In the second step, the TCRαβ–CD3δε bond interacts with CD3γε (kinetic rates *k*_3_ and *k*_–3_) or the TCRαβ–CD3γε bond interacts with CD3δε (kinetic rates *k*_4_ and *k*_–4_) to form CD3γε–TCRαβ–CD3δε trimolecular bonds. Refer to Eq. 2, the modified kinetic differential equations for the steady state are,

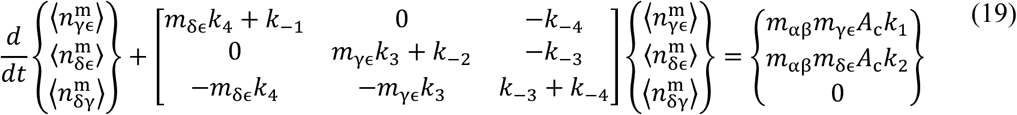

whose solutions are derived as,

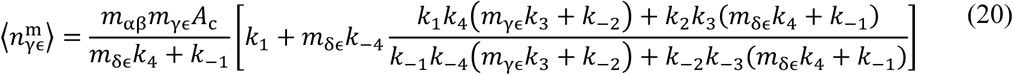

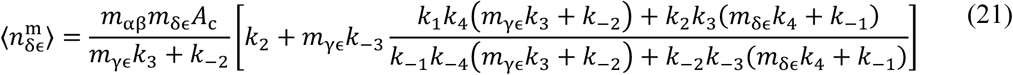

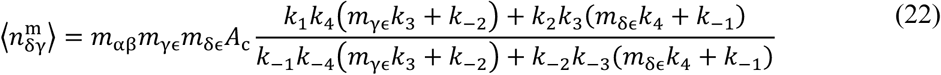

where *m* with different subscriptions indicates the site densities of TCRαβ, CD3γε, and CD3δε, respectively.

Cooperativity arises in the second step by allowing concurrent *cis*-engagements among CD3γε, TCRαβ, and CD3δε. As expected, letting *k*_3_ = *k*_4_ = 0 abolishes the trimolecular bonds and reduces Eqs. 20 and 21 to the forms of Eq. 3 for the TCRαβ–CD3γε and TCRαβ–CD3δε bimolecular bonds. Note that the comparison between the gain in 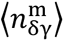 and the loss in ⟨*n*_y**ε**_) – 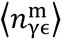 and ⟨*n*δ**ε**⟩ − 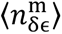 defines the sign of cooperativity. Furthermore, the reduction of bimolecular bond fraction leads to the inequality of 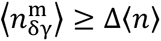. From this, the lower bound for the trimolecular bond fraction can be estimated using the cooperativity value alone as 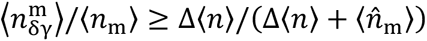(*i.e.*, cooperativity divided by one plus cooperativity). For example, based on the bond number cooperativity of 170% that we measured, the chance of trimolecular bond formation is higher than 62% of the total bonds.

To model the dual-species dissociation processes, similar to the off-rate analysis in the thermal dynamic assay, by setting the right-hand side of Eq. 19 master equations to zero, we obtained the two-step dissociation of dual-species following a hyper-exponential decay. Again, letting the second-step kinetic rates *k*_3_ = *k*_–3_ = *k*_4_ = *k*_–4_ = 0 reduces the master equations to two uncoupled first-order reverse reactions for the TCRαβ–CD3γε and TCRαβ–CD3δε *cis*-dissociation, following single-exponential decays.

### Kinetic modeling of catch-bonds

The *cis*-bond profiles of TCRαβ interacting with CD3γε, CD3δε, or both show catch-slip behavior, with bi-exponential-decay survival probabilities, suggesting two bound species (*98–100*), particularly under clamping force. Therefore, to analyze force-dependent dissociation of bimolecular bonds, we assumed that a single bond might undergo either a fast-dissociation pathway from a weak state or a slow-dissociation pathway from a strong state (indicated by *), and that the two pathways could exchange. To analyze the mixture bond data, in addition to combining the two-pathway models for pairwise TCRαβ–CD3γε and TCRαβ–CD3δε dissociation, we added a third population of trimolecular CD3γε–TCRαβ–CD3δε bonds using the same modeling scheme. The corresponding reaction diagrams for all scenarios are depicted in fig. S2A.

The force dependency of kinetic rates follows Bell’s equation (*101*),

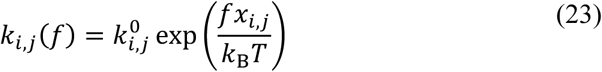

where the subscript *i* denotes rates of dissociation along fast (-f) or slow (-s) pathways and of activation into (a) or deactivation from (-a) the strong states; the subscript *j* denotes different bond types of TCRαβ–CD3γε (γ), TCRαβ–CD3δε (δ), and CD3γε–TCRαβ–CD3δε (δγ); the superscript 0 denotes the respective force-free kinetic rates. According to transition state theory, the force linearly modulates the free energy of the transition barrier, 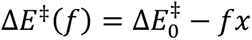, and subsequently regulates the kinetic rates, 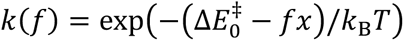, where *x* is the characteristic transition distance, and *k*^0^ is proportional to 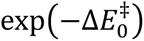.

This two-state, two-pathway dissociation model (*46*) recapitulated the lifetime of catch bonds and the bi-exponential decay of survival probabilities. We obtained the force-free fast dissociation rates for the dimeric bonds from the thermal fluctuation assay, and then evaluated the remaining parameters by globally fitting the model to the force-lifetime distributions via maximum likelihood estimation (MLE). The initial proportions of the two states were determined from the ratios of the kinetic rates, assuming equilibrium. We used a two-step optimization procedure to minimize the MLE: we first explored a broad parameter space using differential evolution to obtain an approximate solution, then refined this using L-BFGS optimization to converge on the minimum. (fig. S2, E to G). The resultant kinetic rates *vs*. force relationships are presented in fig. S2, B to D, and Bell’s parameters are summarized in table S2. We obtained the standard deviation for each parameter by inverting the Hessian matrix at the MLE to approximate the covariance matrix, whose diagonal entries correspond to the variance of the fitted parameters.

### Molecular dynamics simulations

#### 1. Molecular constructs

All simulations were based on the cryo-EM structure (PDB ID: 6JXR) (*28*). The TCRαβ, CD3γε, and CD3δε were truncated near the CPs between extracellular and intracellular regions, leaving the complete TCR–CD3s ECD in its trimeric assembly. The TCRαβ were trimmed at their C-termini, leaving the α-chain Gly22-Glu237 and the β-chain Val26-Glu269. The CD3γε and CD3δε were obtained by truncating the γ-chain after Cys107, ε-chain after Asn121, δ-chain after Glu98, and ε’-chain after Glu124. To minimize potential trimming effects, all chain ends were modified to prevent the exposure of charged residues. An acetyl group was added in front of the N-terminus, and an N-methylamine group was added to cap the C-terminus for each chain. All hydrogen atoms, as well as any missing atoms, were added back through Gromacs utilities (*28*). The disulfide bonds in the original cryo-EM structure were preserved through Gromacs interactive S-S bridge selection procedures (*28*). The resulting CD3γε–TCRαβ–CD3δε trimeric assembly was used to simulate the concurrent *cis*-binding. Then, three partial assemblies were generated to simulate the pairwise *cis*-interactions by removing CD3δε, CD3γε, and TCRαβ individually from the fully assembled ECDs, yielding dimeric TCRαβ–CD3γε, TCRαβ–CD3δε, and CD3γε–CD3δε, respectively.

#### 2. Unit cell preparation

Unit cells enclosing molecular systems of interest were built with physiologically realistic surroundings and then brought into thermodynamically favorable states. The initial structures were oriented and centered within optimized orthorhombic cells, which accommodated the physical extensions in all directions with a ≥ 1.6 nm buffering zone apart from the box sides to ensure the protein was well-isolated. The molecules in unit cells were solvated using the TIP3P water model (*102*). We counterbalanced the negative charges with sodium ions, then set the ionic strength to 150 mM with sodium chloride. To further prepare the system thermodynamically, we eliminated energy-unfavorable close contacts through steepest descent and conjugate gradient energy minimization cycles. While restraining the heavy atoms in macromolecules, we brought the unit cell to a suitable ensemble using a 200-ps velocity-rescaling simulation to adjust the temperature and a 200-ps Berendsen Barostat simulation to correct the pressure (300 K, 1.01325 bar). After releasing the heavy-atom restraints, the systems were considered ready for subsequent equilibration and production runs.

#### 3. Conventional molecular dynamics

CMD simulations were performed using GROMACS (version 2019.6) (*103–105*) under the recent AMBER99SB*-ILDNP force field with improved helix-coil transitional and side-chain torsional dynamics (*106*). From previously prepared unit cells, we randomly sampled the initial velocities from the Maxwell-Boltzmann distribution. We self-evolved the molecular trajectories using the leap-frog algorithm, integrating Newton’s law of motion every 2 fs. For CMD, the individual simulation runs were executed under periodic boundary conditions to sample the NVT ensemble under corresponding constant volumes (*i.e.*, CD3γε–TCRαβ–CD3δε: 12.5 × 10.8 × 9.9 nm^3^; TCR–CD3γε: 12.6 × 9.8 × 8.6 nm^3^; TCR–CD3δε: 11.9 × 10.3 × 9.2 nm^3^; CD3γε–CD3δε: 11.5 × 8.1 × 8.0 nm^3^) and 300 K constant temperature. The neighborhood lists were refreshed every 10 steps using the buffered Verlet scheme, with a 1.2 nm non-bonded cutoff for Coulomb and van der Waals interactions. We computed long-range electrostatic potentials using the Particle Ewald Mesh summation method. All systems were simulated for 200 ns with three replicates, with each trajectory including at least a 50 ns equilibration stage and the remaining production run. Conformational snapshots every 0.1 ns during the production stage were extracted for subsequent analysis.

#### 4. Steered molecular dynamics

SMD simulations were initiated from the well-equilibrated configurations from corresponding CMD simulations (*i.e.*, the conformational snapshots at 100 ns). We prepared them according to the Unit cell preparation section, but with a 20-nm box length to accommodate molecular extension and separation under external pulling. The external forces with constant magnitudes (*i.e.*, force-clamp) or constant velocities (*i.e.*, force-ramp) along the fixed *z*-direction were added to the trimeric and dimeric complexes. We applied the clamping or ramping forces to the N-terminus of the TCRα chain to accelerate dissociation while keeping the C-termini of the CD3ε chains fixed to mimic the anchor effect of their TM domains. For force-clamp simulations, we generated five independent trajectories at 100, 125, 150, 175, and 200 pN for each of the two dimeric systems, but we ran only three independent runs at 150, 175, and 200 pN for the trimeric complex. We aborted the CD3γε–TCRαβ–CD3δε simulations at smaller clamp forces of 100 and 125 pN due to the inability to observe dissociation events after the exceedingly long waiting time (> 1000 ns). For force-ramp simulations, a 70 pN/nm pseudo spring was utilized to exert increasing levels of pulling to accelerate dissociations. One end of the spring was linked to the α chain N-terminus of the targeting molecule, and the other free end departed in the *z*-direction at 1 nm/ns. We performed five repeats each for two dimeric and one trimeric ECD complexes to assess corresponding rupture forces as a consistency check.

### Occupancy analyses

We used several variations of occupancy analyses to dynamically evaluate the extent to which a residue remains in contact with its binding partner throughout the simulation. At timestamp *i*, the residue being analyzed is defined to be in contact (denoted *o_i_* = 1) if the center-of-mass of this residue is within 4 Å of any atom of its binding partner; otherwise, it is not in contact (denoted *o_i_* = 0). The occupancy index (%) indicates the fraction of time in contact during a given period, as ∑*_i_ o_i_* /*N*, where *N* is the total number of analyzed frames.

#### 1. Residue-to-residue occupancy (fig. S5)

By setting residues within a subunit as a reference group, this occupancy analysis quantifies the extent of dynamic interaction between each residue in one subunit and every residue in the other over a given period. Thus, it evaluates the level of any pairwise contacts, thereby forming a correlation matrix where the column indexes the residues from one subunit; the row indexes the residues from the other subunit, and the elements denote the values of residue-to-residue relative binding strength, ranging from 0 to 1.

#### 2. Residue-to-domain occupancy (fig. S6)

This occupancy analysis quantifies each residue against all residues within its binding partner, using the entire subunit as the reference group. It is usually smaller than the sum of residue-to-residue occupancy over the binding domain because the residue in question may simultaneously contact multiple residues in the domain under consideration; however, these contacts can be counted only once for that particular residue. To broaden its dynamic range, ΣOccupancy is defined as the integration of the CMD and SMD row-/column-wise sums of residue-to-residue occupancies, while ΔOccupancy is defined as the differentiation of the CMD from the SMD row-/column-wise sums of residue-to-residue occupancies, which were calculated to summarize and discriminate force-free and force-clamp interface dynamics (Fig. 4, G and H).

#### 3. Domain-to-domain occupancy

This occupancy analysis further reduces data dimensionality to examine overall binding tendency against the other domain by averaging over all interacting residues (*i.e.,* positive terms of residue-to-domain occupancies), providing a measure of overall domain-to-domain adhesion strength.

### Linear regression

We conducted linear regression to assess correlations between *cis*-, *trans*-kinetic measures (*x*) and functional outputs (*y*), between *cis*- (*x*) and *trans*-kinetics (*y*), or between MTP readouts (*x*) and IL-2 (*y*). The goodness-of-fit was quantified using Pearson’s correlation coefficient, *i.e.*, 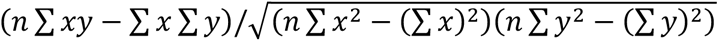, which preserves the sign of correlation, as well as its square, *R*^2^. The sensitivity-of-prediction, *i.e.*, 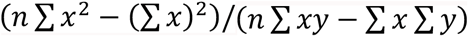, is the inverse of the linear-regression slope.

### Molecular tension probe assay

DNA-based MTPs were prepared as described (*78, 107*). Briefly, an MTP is generated by thermal annealing three single-stranded (ss) DNAs in high-salt buffer on a glass surface prepared through sequential treatments: piranha oxidation, salinization with 2% 3-aminopropyltriethoxysilane in ethanol, and surface blocking with BSA. The middle portion of one ssDNA was folded into a force-sensitive hairpin, with its two end portions hybridized to two other ssDNAs to form two duplexes that position a fluorophore and quencher pair in proximity when the hairpin is folded. The other ends of the DNA duplexes were attached to a glass surface and a pMHC/BSA, respectively.

### Spread and force imaging

Jurkat cells expressing 2B4 TCR constructs were plated on glass surface functionalized with MTP tagged with K5:I-E^k^ or BSA (10 µg/ml) in imaging buffer (1× HBSS supplemented with 10 mM HEPES) containing a locker ssDNA (1 µM) complementary to the unfolded DNA hairpin and allowed to adhere for approximately 10 min before image acquisition, which was performed on a Nikon W1 spinning-disk confocal system equipped with a Plan-Apochromat 60×/1.40 NA oil-immersion objective. We employed three imaging modalities simultaneously: differential interference contrast (DIC) microscopy to provide a top view of cells, reflective interference contrast microscopy (RICM) to monitor cell-substrate contact morphology, and total internal reflection fluorescence (TIRF) microscopy to detect force-dependent fluorophore dequenching from unfolded hairpin constructs. This tri-modal approach enabled simultaneous measurements of cellular spreading area and number of opened MTPs across ligand-receptor interfaces.

### Statistical assessments between bond profiles

To test whether two conditions (*e.g.*, WT *vs.* mutant) have significantly different force-lifetime relationships, we fitted bond lifetime versus force data using a two-state catch-slip bond model with eight parameters per curve (fig. S2). We compared a full model (separate curves for each condition) to a reduced model (single pooled curve for both conditions) using a likelihood ratio test (LRT): **Λ** = 2(log ℒ_full_ − log ℒ_reduced_) ∼ *χ*^2^(df), with degrees of freedom (df) equal to the difference in parameters between the full and reduced models. A significant LRT statistic (**Λ**) indicates that the two conditions have significantly different force-lifetime curves.

To test statistical significance between bond profiles at individual force levels, we used *z*-tests to assess error overlaps between the mean bond lifetimes of the two conditions. The force-dependent lifetime generally follows a hyper-exponential distribution dominated by a single component (as *p*_1_ ≫ *p*_2_, *μ*_1_ ≫ *μ*_2_), which can be approximated as a single exponential with a standard deviation estimated from its mean. For each force level, the mean (*μ*) and standard deviation (*σ*) of an exponential distribution are both equal to the reciprocal of the rate parameter *λ*: *μ* = *σ* = 1/*λ*. The *z*-score for comparing two means is calculated as, 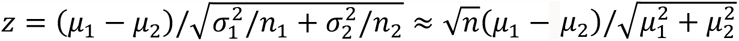, where *n* denotes the number of measurements per force bin.

## ACKNOWLEDGEMENTS

This work was supported by NIH grants R01GM124489 (to M.K.) and R01CA284604 (to M.K. and C.Z.) and U01CA214354 (to C.Z. and M.K.). The MD simulations were supported by an NSF award MCA08X014 (to C.Z.) in advanced computing infrastructure for the U.S. and were performed in the Extreme Science and Engineering Discovery Environment (XSEDE). Cell sorting/flow cytometry technologies were provided by NYU Langone’s Cytometry and Cell Sorting Laboratory, which is supported in part by NIH grant P30CA016087. I-E^k^ pMHC was made by the NIH Tetramer Core Facility at Emory University. MTP experiment was supported by Center on Probes for Molecular Mechanotechnology at Emory University, which is funded by NIH grant RM1GM145394. We thank Larissa Doudy and Courtney Ng for technical assistance, Johannes Huppa for providing the H57 scFv, and Bernard Malissen for providing mouse 58^−/-^ T cell hybridoma cells.

## AUTHOR CONTRIBUTIONS

C.Z., P.C., and C.G. conceived the study; C.Z. and M.K. directed the study; P.C., Z.Y., C.G., R.D. and A.N. performed experiments; P.C. performed simulations; P.C. and A.N. performed structural analysis; A.N., S.B., X.L, and D.G. produced proteins; P.C., S.T. and C.Z. constructed mathematical models and analyzed data using the models; J.S. drew the model illustration. P.C., A.N., Z.Y., C.G., R.D., M.K., and C.Z. analyzed data and wrote the paper with contributions from other authors.

