## Supplementary materials for "Cooperative *cis*-interactions between ectodomains of TCRαβ and CD3 subunits enable mechanotransduction"

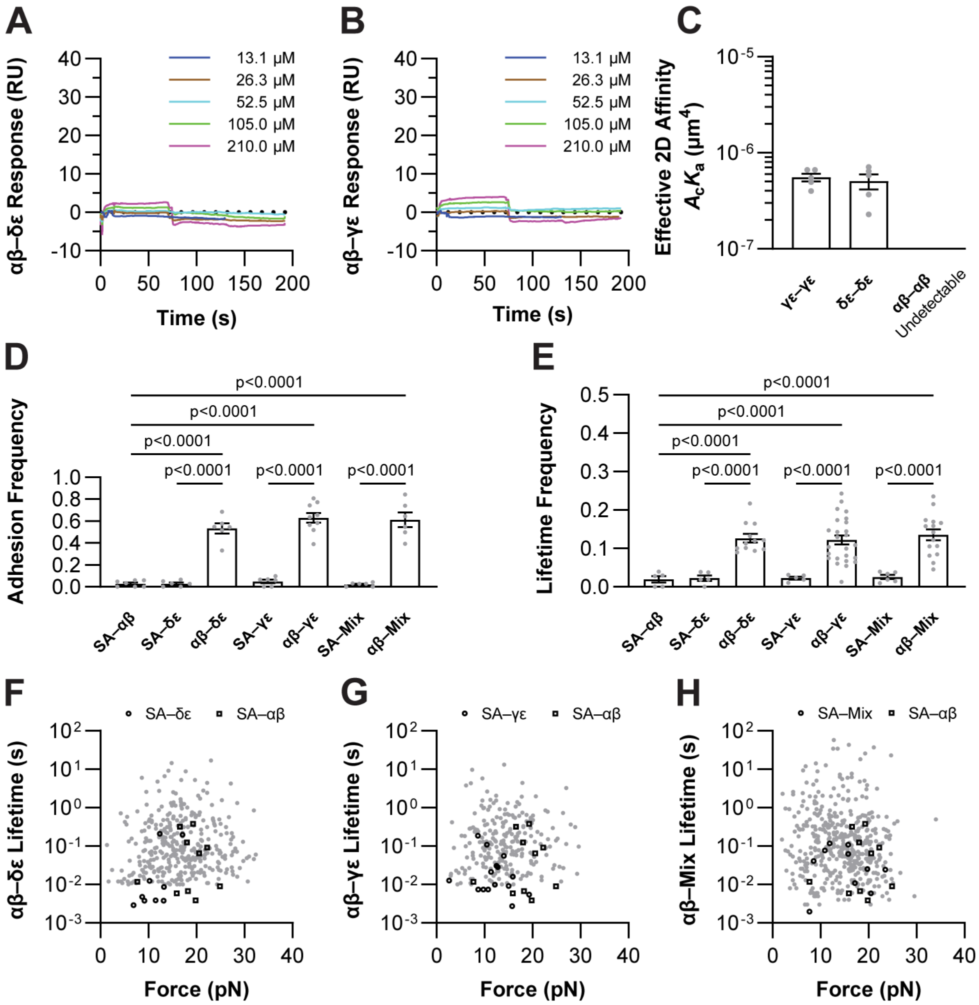

**Figure S1.** (**A** and **B**) **Inability of SPR to detect TCR–CD3 3D binding.** SPR measurements were attempted using the BiaCore T200 instrument. 10 μg/mL biotinylated CD3γε and CD3δε were immobilized on streptavidin (SA) coated sensor chips. Different concentrations of analyte solution with 13.1, 26.3, 52.5, 105, and 210 μM (*colored lines*) TCRαβ were injected over surface-bound CD3γε (A) and CD3δε (B) at time 0, which was then switched to buffer at 75 s. The recorded SPR signals were processed using T200 evaluation software 3.0, but it failed to obtain any reliable kinetic parameters. (**C**) **Effective 2D affinities of homotypic interactions of CD3γε, CD3δε, and TCRαβ ECDs.** The homotypic 2D affinities were measured similarly to the heterotypic 2D affinities described in the main text by the MAF assay, but with significantly lower values. Note that the 2B4 TCRαβ homotypic interactions were undetectable using current experimental setups. (**D**) **Specificity controls for the MAF assay.** Multiple negative controls, where one reactant was omitted, were tested to ensure the specificity of our TCRαβ–CD3γε, TCRαβ–CD3δε, and TCRαβ–CD3m measurements. SA here indicates RBCs without further attachments of biotinylated reactants. The background-level adhesion frequencies were observed when either binding partner was missing, while significantly higher adhesion frequencies were observed when both binding partners were present. (**E**) **Specificity controls for the BFP force-clamp and thermal-**

**fluctuation assays.** Similar negative controls were conducted for our TCRαβ–CD3γε, TCRαβ–CD3δε, and TCRαβ–CD3m bond lifetime measures by omitting one reactant from the interacting pair. The apparent differences between experiments and controls validate the relevance of our BFP signals. Lifetime frequency is the fraction of lifetime events in the total adhesion events. Note that the adhesion frequency in BFP experiment was intentionally kept at $\leq$ 20% to minimize the probability of multi-bond formation. Data in (D) and (E) are presented as mean ± SEM of individual measurements. Statistics were obtained by comparing any two groups using unpaired Student's t-tests. (**F** to **H**) **Signal-to-noise comparison for BFP assays.** The bindings in negative controls occasionally resulted in bond lifetime events as well. These non-specific bond lifetimes (*open symbols*) are not only less frequent but also shorter-lived when compared against those from TCRαβ–CD3γε (F), TCRαβ–CD3δε (G), and TCRαβ–CD3m (H) bond lifetimes (*closed symbols*).

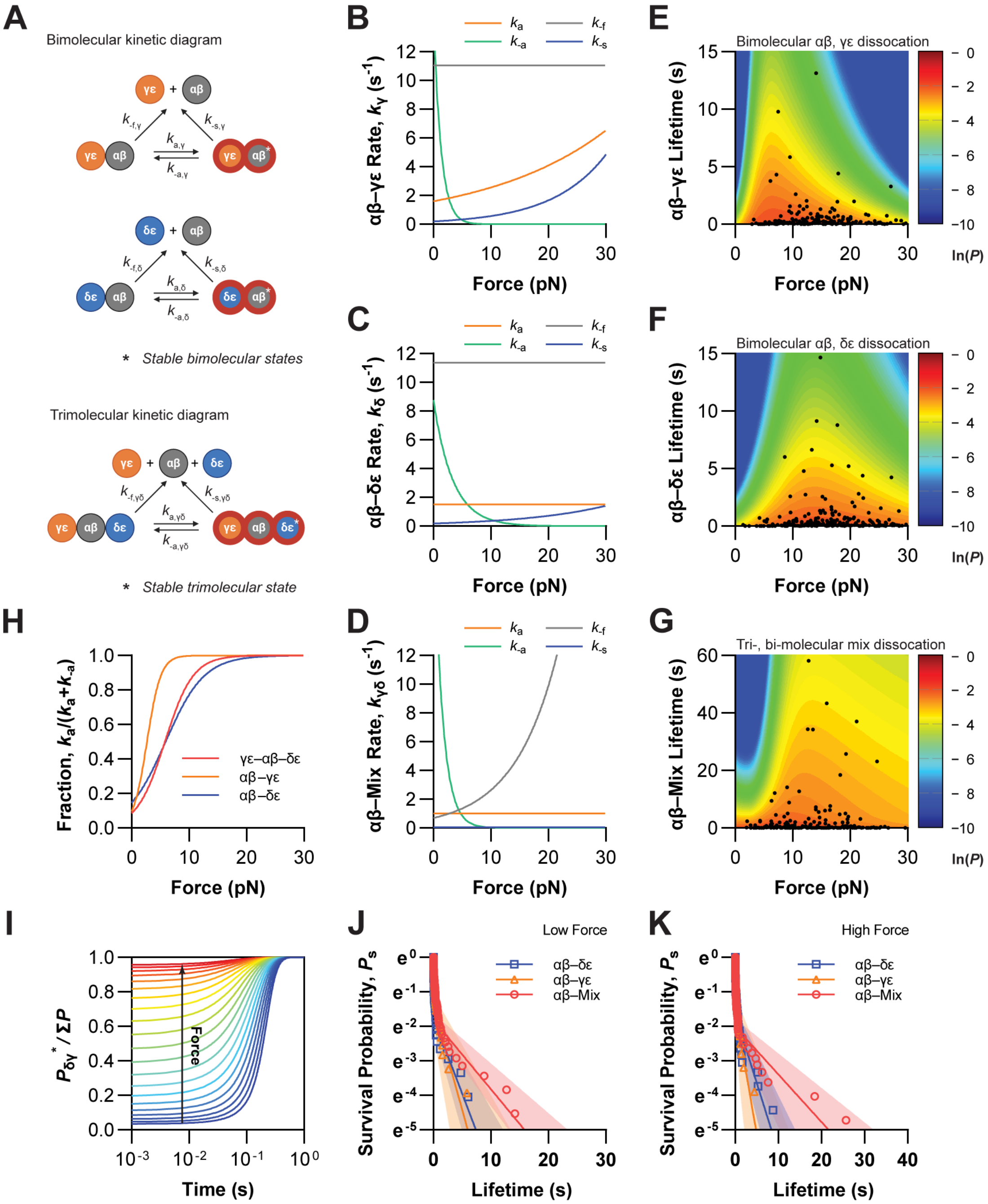

**Figure S2. (A) Kinetic mechanisms of the two-state, two-pathway dissociation models for catch-slip bonds.** The schematics for TCRαβ–CD3γε (*top*) and TCRαβ–CD3δε (*middle*) bimolecular interactions depict the two kinetic pathways dissociating from two interconverting bound states, with arrows indicating possible reaction directions and symbols denoting corresponding reaction rates. For the case of TCRαβ–CD3m, the CD3δε–TCRαβ–CD3γε trimolecular bound states (*bottom*) are attainable, in addition to the TCRαβ–CD3γε and TCRαβ–CD3δε bimolecular bound states. Note that the superscripts * on the reactant label the strong bound states. These kinetic mechanisms require a total of 12 rate coefficients ($k_{-\mathrm{f}}$, $k_{-\mathrm{s}}$, and $k_{\pm\mathrm{a}}$ for γ, δ, and γδ), each following Bell's model (see Methods). The subscripts a and -a indicate activation and deactivation between the weak and strong bound states, -f and -s indicate dissociations along the fast and slow pathways, and the Greek letters label the types of reactants. (**B** to **D**) **Reaction rates as a function of force.** After fitting the experimentally obtained bond lifetime, we derived the best-fit Bell parameter sets for the TCRαβ–CD3γε (B), TCRαβ–CD3δε (C), and TCRαβ–CD3m (D) interactions, as listed in table S2. Based on Eq. 23, the corresponding kinetic rates are plotted *vs.* force for the fast and slow dissociations, as well as the activation and deactivation transitions between the two states. (**E** to **G**) **Bond lifetime scatter plots fitted with the catch-slip bonds model using maximum likelihood estimation.** The probability of observing the data (*points*) under the model prediction (*curves*) was maximized to obtain the best-fit parameter sets. The contours indicate the likelihoods, $\ln(P)$ *vs.* time and force for the TCRαβ–CD3γε (E), TCRαβ–CD3δε (F), and TCRαβ–CD3m (G) interactions. All model parameters were iteratively derived from global fitting, except for the fast dissociation rates, $k_{-\mathrm{f}}^{0}$, which were pre-determined from the BFP thermal fluctuation assay (Fig. 2E). The model-predicted mean ± SD curves well capture the average bond lifetime from simple force binning (*points with error bars*) in Fig. 2D. (**H**) **The effect of force on reaction states for CD3δε–TCRαβ–CD3γε.** The fractions of strong binding states, $k_{\mathrm{a}}/(k_{\mathrm{a}} + k_{-\mathrm{a}})$, as evaluated using activation and deactivation transition kinetics for the two bimolecular and one trimolecular interaction, are plotted against force. (**I**) **The effect of force and time on strong binding states for CD3δε–TCRαβ–CD3γε.** The probabilities of long-lasting, strong binding states are plotted as a function of time and force. (**J** and **K**) **The bond survival probability under force,** derived from BFP bond lifetimes subject to 10 pN (J) and 20 pN (K) tensile forces for TCRαβ–CD3m (*red*) in comparison with TCRαβ–CD3γε (*orange*), TCRαβ–CD3δε (*blue*) interactions.

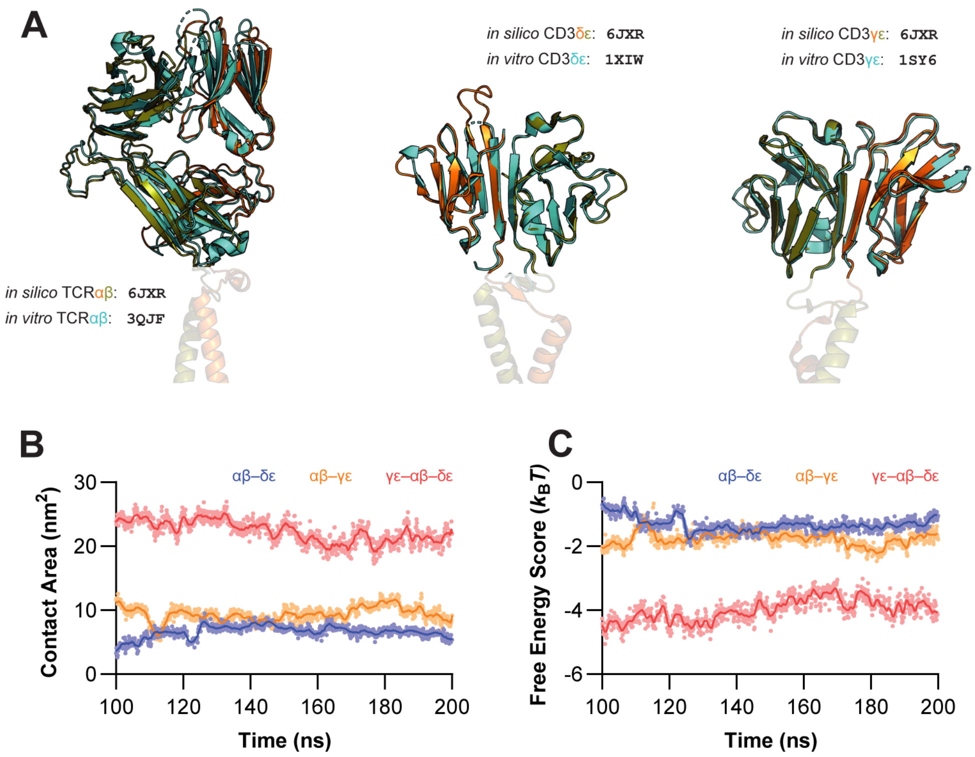

**Figure S3.** (**A**) **Structural comparisons between *in vitro* and *in silico* investigations.** The similarities between experiments and simulations are compared by superimposing structures for the TCRαβ (*left*), CD3γε (*middle*), and CD3δε (*right*) ECDs resolved using cryo-EM (*orange and olive*, PDB: 6JXR) (*28*) and crystallography (*cyan*, 3QJF, 1SY6, and 1XIW, respectively) (*63, 94, 110*). The recent cryo-EM co-complex revealed the tetrameric arrangement of TCR–CD3 complexes, which served as the initial models for our MD simulation. On the other hand, the crystal structures were based on the same molecular constructs as the individual hybrid 2B4 TCR and human CD3 proteins used in *in vitro* experiments. (**B** and **C**) **Contact areas and interaction energy dynamics.** The full 100 ns dynamics trace the contact areas (B) and free energy scores (C) between TCRαβ and CD3δε (*blue*), CD3γε (*orange*), and both (*red*), where the molecular structures per 100 ps from 100 to 200 ns equilibrated CMD trajectories were analyzed using Gromacs 2019 sasa tool (*104*) and Autodock Vina 1.1.2 (*111*), respectively.

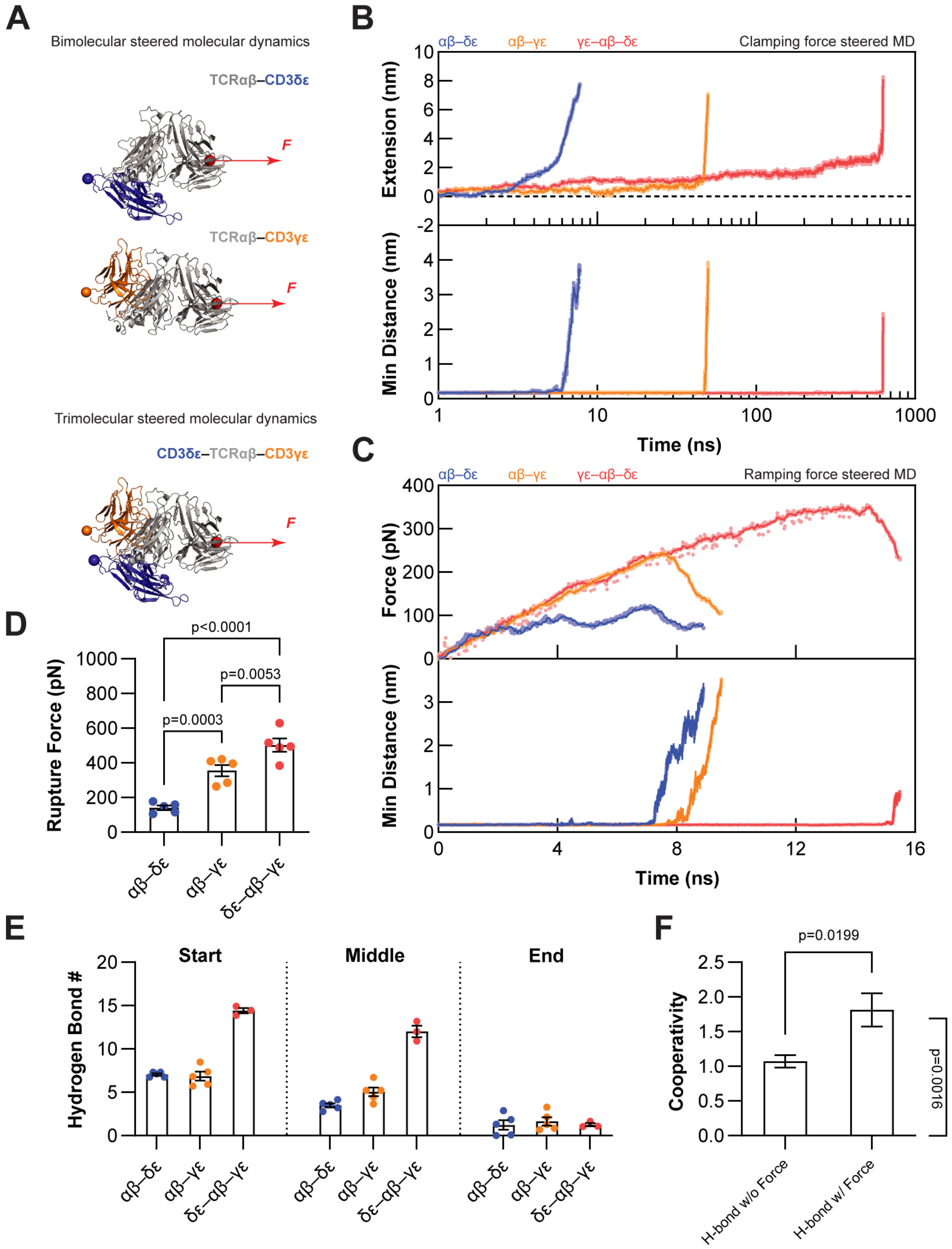

**Figure S4.** (**A**) **Initial structures and pulling geometries for SMD simulations.** Initial structures for SMD simulations were generated by extracting equilibrated bimolecular and trimolecular constructs from the middle of the CMD simulations. Forces (*red*) were applied to the TCRα N-terminus, pulling away from the anchored C-termini of CD3γε (*top*), CD3δε (*middle*), and both (*bottom*). Two types of force applications were implemented: force-clamp and force-ramp SMD simulations. For the former, the pulling force was maintained at a constant magnitude; for the latter, it was increased stepwise from zero with umbrella pulling. (**B**) **Representative trajectories of**

**extensions and min-distances of force-clamp simulations.** The extensions measuring the end-to-end distances (*top*) of TCR–CD3s reflect the overall elongation of the molecule under pulling. Abrupt increases in extension are commonly associated with significant structural changes within or between proteins, such as dissociations. Correlatively, the minimum distances (*bottom*) between TCRαβ and CD3s follow similar trends but precisely pinpoint the onsets of dissociation events. The min-distance trajectories shown here are from different simulations than those in Fig. 2G, but the times to dissociation remain the same order as δε–αβ < γε–αβ < δε–αβ–γε. (**C**) **Representative trajectories of forces and min-distances of force-ramp simulations.** In force-ramp simulations, the molecular extensions increase over time, but the force levels can increase or decrease depending on the speed of pulling relative to that of the molecular extension (*top*). Structural failures within or between proteins lead to their abrupt elongations, resulting in sudden drops in force magnitude. Again, high correlations are observed when compared with the minimum-distance traces (*bottom*). The maximum forces and the forces at molecular separation, *i.e.*, the rupture forces, as alternative readouts of binding stability, show the same rank order: δε–αβ < γε–αβ < δε–αβ–γε. (**D**) **Rupture forces of the TCR–CD3 interactions.** Five repeats of force-ramp SMD simulations observed similar rupture-force measures, confirming the rankings in (C) and (B). The p-values were calculated by unpaired Student's t-tests. (**E**) **H-bonds during force-clamp simulations.** We dissected the TCR–CD3 interactions by focusing on the H-bond dynamics, given their crucial role in non-covalent interactions. An H-bond was identified by a distance < 3.5 Å between donor and acceptor atoms and an angle of $180^{\circ} \pm 30^{\circ}$ between them. The instantaneous H-bond counts between TCRαβ and CD3γε, CD3δε, and both were tracked throughout SMD simulations. H-bond counts are summarized using the mean of total H-bond numbers in the start (first 1 ns, *left*), middle (remaining, *middle*), and end (last 1 ns, *right*) periods, where each point represents average H-bond number of individual SMD runs under 100, 125, 150, 175, and 200 pN forces for dimeric interactions, and 150, 175, and 200 pN forces for trimeric interactions, respectively. The H-bonds were identified using the Gromacs 2019 hbond program (*104*). (**F**) **Cooperative analysis of the hydrogen bond (H-bond).** A similar cooperative analysis reports positive cooperativity both without and with force in terms of H-bond counts. Again, stronger cooperativity is observed in the presence of force. Statistical significance of the force effect and the cooperativity were assessed, respectively, by unpaired Student's t-tests and one-sample t-tests against zero.

A

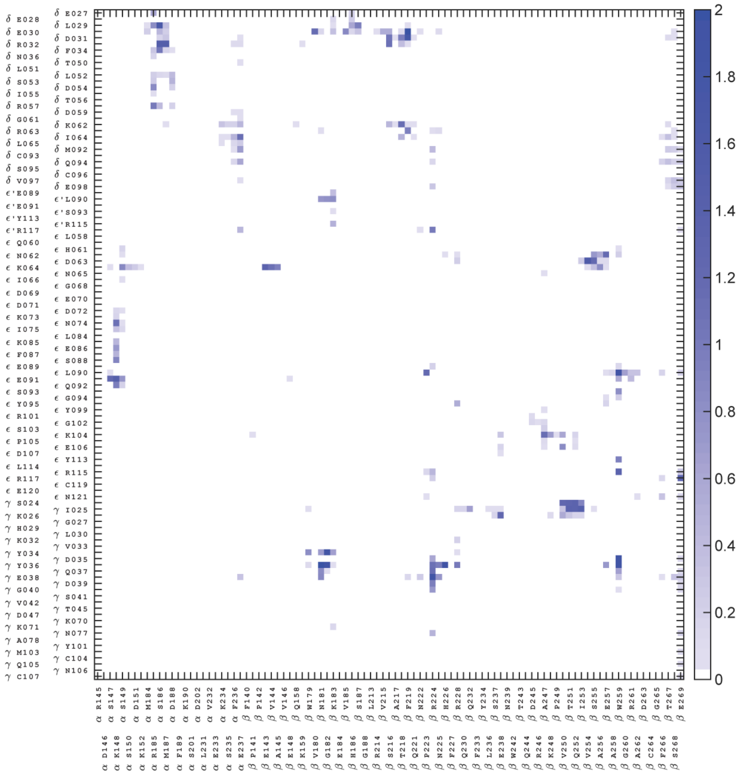

B

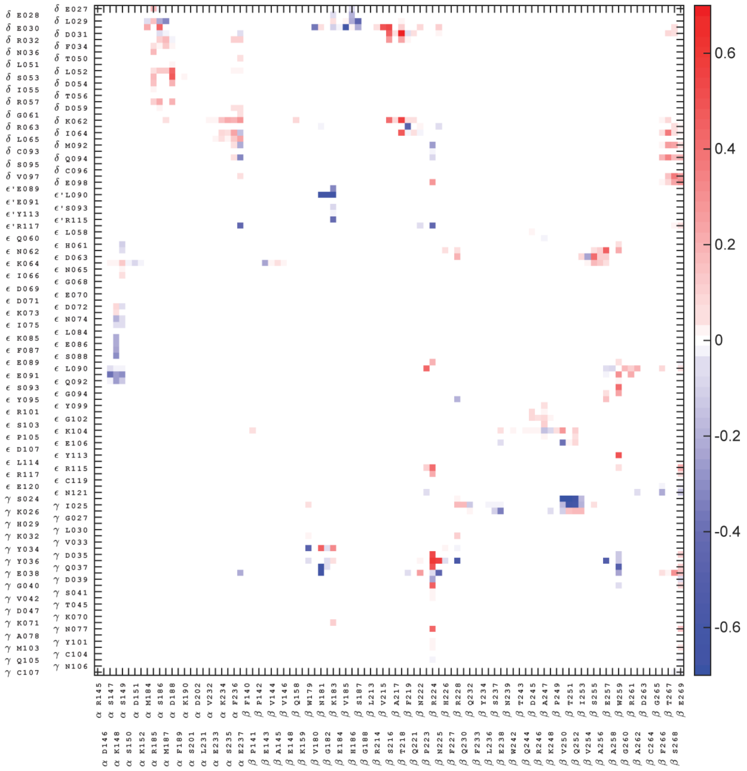

**Figure S5. Residue-to-residue occupancy matrices of the TCR–CD3 interfaces.** (**A** and **B**) Matrices display the residue-to-residue occupancy values for the CD3δε–TCRαβ–CD3γε trimeric complex, where each value denotes the level of interaction between the TCRαβ residue (*column index*) and the CD3s residue (*row index*) (see Methods). TCR residue indices (*bottom*) correspond to TCR α and β chain amino acid (a.a.) aligned from N to C-termini. CD3 residue indices (*left*) stand for a.a. of CD3δ, ε', ε, and γ chains aligned from N- to C-termini. The residue name begins with the chain name, followed by the a.a. abbreviation and residue number. Here, the zero entries of the matrices have been removed for clarity, leaving only TCR–CD3 interacting residues. Two versions are shown: (A) a Summation matrix, combining CMD and SMD occupancies element-by-element; (B) a Difference matrix, subtracting CMD occupancies from SMD. The level of color saturation in (A) correlates with the level of interactions, gradually transitioning from *white* to *blue* as the summation value increases from 0 to 2. The color coding in (B) denotes the force-induced changes, using *blue* to *white* to *red* to present negative to zero to positive difference values. So, *red* indicates mechano-enhanced and *blue* signifies mechano-suppressed residue-to-residue contacts.

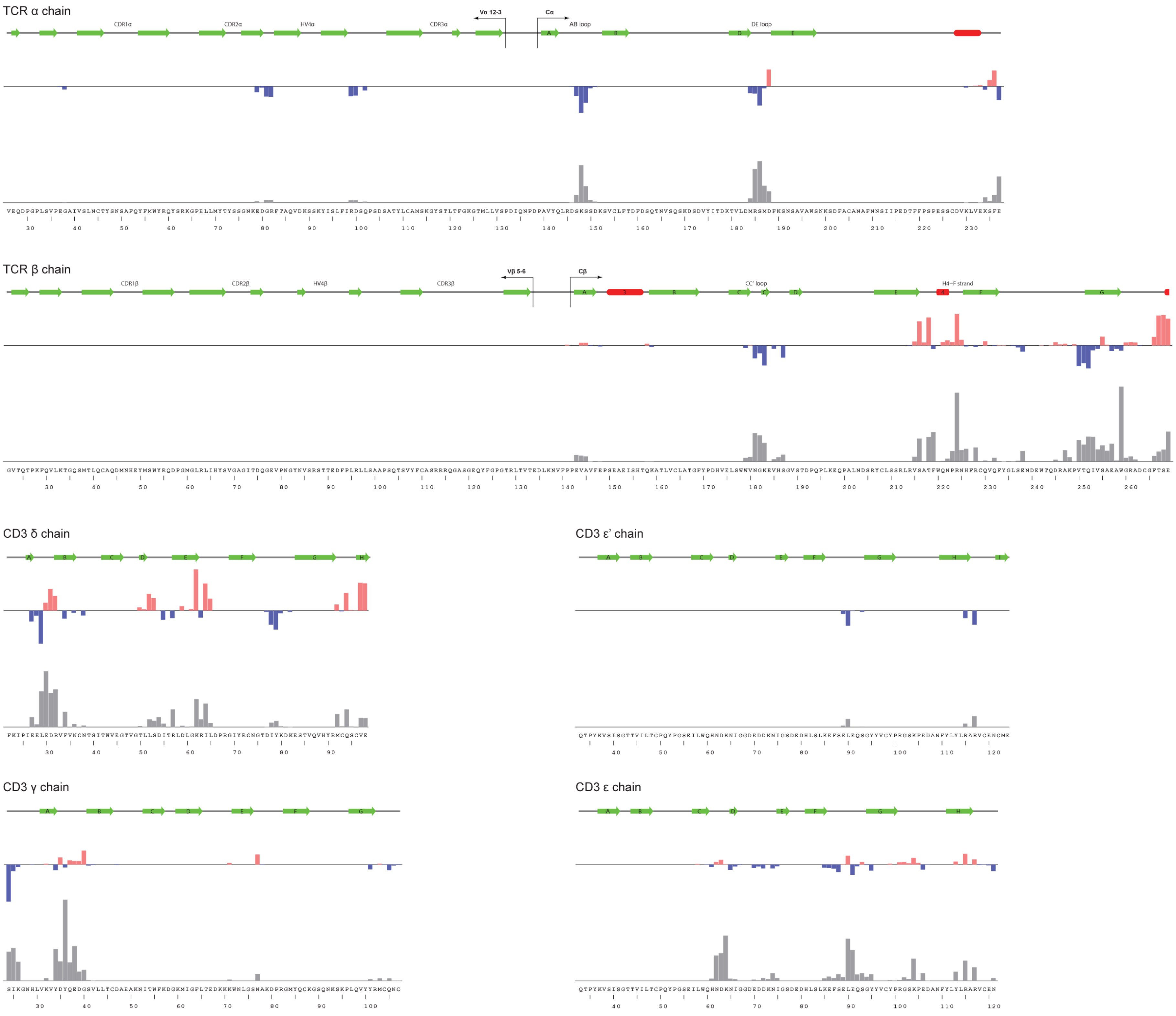
TCR α chain
CDR1α
CDR2α
HV4α
CDR3α
Vα 12-3
Cα
AB loop
DE loop
TCR β chain
CDR1β
CDR2β
HV4β
CDR3β
Vβ 5-6
Cβ
CC' loop
H4–F strand
CD3 δ chain
CD3 ε' chain
CD3 γ chain
CD3 ε chain

**Figure S6. Residue-to-domain occupancy profiles of the CD3δε–TCRαβ–CD3γε trimeric complex.** Bar graphs display the residue-to-domain occupancy values for TCRαβ and CD3s, by summing over the matrices in fig. S5 by column and by row, respectively (see Methods). The residue-to-domain profiles for TCR α, β, and CD3δ, ε', ε, and γ chains are presented using vertical bars, where the height denotes the level of interactions of the single TCR residue with all CD3s or the particular CD3 residue with the entire TCRαβ. The residue names and indices are listed at the *bottom*, while the corresponding secondary structures are displayed at the *top* (β-strand in *green* and α-helix in *red*). Note that interactions are usually at loops between well-defined secondary structures, and the text in the structural diagram highlights the regions of interest, such as the AB loop. Similar to fig. S5, two versions of occupancies are presented here: summative ΣOccupancy in *gray*, representing the overall interacting levels, obtained from the CMD and SMD occupancies; and differential ΔOccupancy in *red* and *blue*, obtained by subtracting the CMD from the SMD occupancies, denoting the changes upon force application. So, *red* features force-enhancement, and *blue* marks force-suppression on the residue-domain contacts. The *bars* for two profiles are drawn on the same scale for different chains.

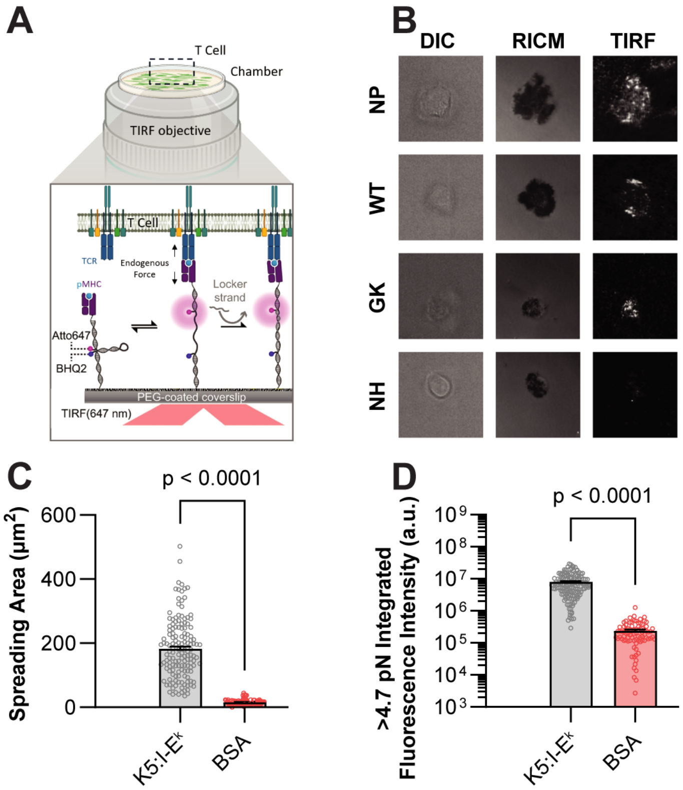

**Figure S7. Mechanosignaling of TCRαβ Variant T Cells** (**A**) **Schematic of the molecular tension probe (MTP) assay.** A DNA-based assembly containing a force-sensitive hairpin flanked by a fluorophore and quencher was immobilized on the glass surface, with K5:I-$E^k$ pMHC (10 μg/ml) or BSA (for negative control) presented at the free end (*left*) for binding by 2B4 TCR expressed on Jurkat cells. TCR–pMHC engagement triggers mechanosignaling for the cell to activate its intracellular force generation and transmission machinery to exert endogenous force on the TCR–pMHC bonds. Above threshold forces (4.7 pN) unfold the hairpin to separate the quencher and the fluorophore to generate a fluorescent signal (*middle*). A complementary DNA strand in solution (1 μM) then hybridize the open hairpin to lock it in the unfolded configuration, which allows the fluorescent signal to accumulate (*right*). (**B**) **Representative MTP images** showing a Jurkat cell expressing NP (*1$^{st}$ row*), WT (*2$^{nd}$ row*), GK (*3$^{rd}$ row*), or NH (*4$^{th}$ row*) 2B4 TCR. Cell morphology (DIC, *left column*), cell spreading (RICM, *middle column*), and TCR tension (TIRF, *right column*) were visualized simultaneously for a single cell. (**C** and **D**) **MTP negative controls.** Mean ± SEM of spreading areas (C) and integrated fluorescence intensities (D) of individual cells (*points*) transduced with WT 2B4 TCR reacting with K5:I-$E^k$ (*gray*) or BSA (*red*) tagged MTP, which ensure the MTP signals came from TCR–pMHC-specific antigen recognition.

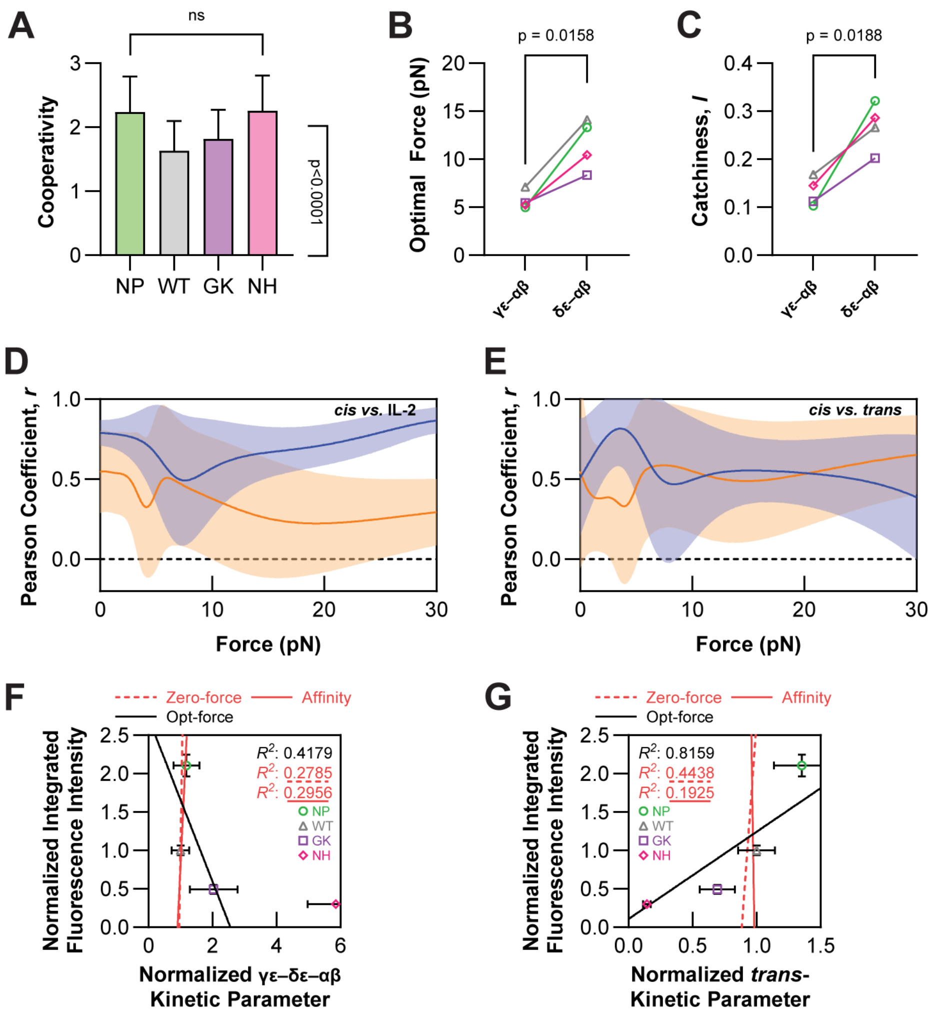

**Figure S8.** (**A**) **Cooperativity analysis of the β-chain double-mutant TCRs.** Cooperativity metrics in terms of average number of bonds, $\Delta\langle n_\mathrm{m}\rangle/\langle\hat{n}_\mathrm{m}\rangle$, as in Fig. 2C, were calculated for CD3δε–CD3γε *cis*-interaction with TCRαβ WT (*gray*) or β-chain double-mutants NP (*green*), GK (*purple*), NH (*magenta*) based on data in Fig. 5G. Statistical significance of mutational effects was assessed by an unpaired Student's t-test, which is not significant (ns), and the cooperativity metrics themselves were assessed by a one-sample t-test against zero, which is highly significant. (**B** and **C**) **Asymmetry of CD3γε and CD3δε *cis*-bonds with TCRαβ** WT and mutants**.** Optimal forces (B) and catchiness (C) of CD3γε (*left*) and CD3δε (*right*) *cis*-catch bonds with TCRαβ WT (*gray*) and NP (*green*), GK (*purple*), or NH (*magenta*) mutants are compared. Data points for the same TCR construct are connected by a line. (**D**) **Correlation between the γε–αβ and δε–αβ *cis*-bond profiles and T-cell function.** The Pearson correlation coefficients of the bimolecular *cis*-bond profiles of WT and MT TCRαβs with CD3γε (*orange*, from Fig. 5H) or CD3δε (*blue*, from Fig. 5I) *vs.* IL-2 production by pMHC-stimulated hybridoma cells expressing the same TCRs (from Fig. 5C) are shown across all forces (*curves with shades*, mean ± SD). (**E**) **Correlation between the**

**γε–αβ and δε–αβ *cis*-bond profiles and the TCR–pMHC *trans*-bond profiles.** The Pearson correlation coefficients of the bimolecular *cis*-bond lifetimes of CD3γε (*orange*, from Fig. 5H) or CD3δε (*blue*, from Fig. 5I) *vs. trans*-bond lifetimes of pMHC with cell surface WT and MT TCRs (from Fig. 5K) are shown across all forces (*curves with shades*, mean ± SD). (**F**) **Inverse correlation between force signals and γε–αβ–δε *cis*-interaction in the presence (but not absence) of force.** The mean ± SEMs of the normalized integrated fluorescence intensities (Fig. 5E) are plotted against the peak lifetimes of the γε–αβ–δε *cis*-bonds (Fig. 5J). The poor correlations and sensitivities of the force signal with zero-force bond lifetime (*red dashed line*) or normalized number of bonds (*red solid line*) of the γε–αβ–δε *cis*-interactions are plotted for comparison. (**G**) **Correlation between force signals and TCR–pMHC *trans*-interactions in the presence (but not absence) of force.** The mean ± SEMs of the normalized integrated fluorescence intensities (Fig. 5E) are plotted against the peak lifetimes of the TCR–pMHC *trans*-bonds (Fig. 5K). The poor correlations and sensitivities of the force signal with zero-force bond lifetime (*red dashed line*) or normalized number of bonds (*red solid line*) of the TCR–pMHC *trans*-interactions are plotted for comparison.

### SUPPLEMENTARY TEXT

By introducing a novel occupancy analysis of domain-binding interfaces, we identified the key *cis*-interacting residues of TCR–CD3 ECDs. These high-occupancy residues correlate well with previously reported mutagenesis (table S3) and affect subdomain assembly and effector functions (*25, 26, 28*). For example, the Cβ CC' loop interacts with the CD3γ AB loop (*28*) through βG182–γY34 (89% occupancy) and βG182–γY36 (96%). Substitutions at βG182R suppressed TCR surface expression (*22*), while βG182A and βG182H mutations reduced cytokine production upon stimulation in Jurkat and T cell hybridoma assays (*26, 27*). Consistent with NMR chemical perturbation experiments (*25, 26*), we also observed interactions between Cβ helix 4 and CD3δ AB and EF loops through hydrophobic contacts (fig. S5) (*28*). Cβ helix 4 mutations βT218R and βF219R, as well as substitutions within the CD3δ AB loop, reduced TCR surface expression by 70-80% (*22*). A nearby βS216R mutation diminished cytokine and CD69 induction upon stimulation (*27*). Similarly, the Cβ helix 4–F strand is a hot spot for CD3γ, δ, and ε chain associations, where residues βR224, βN225 (*25, 26*), and βH226 (*28*) are critical. Correspondingly, the βR224E and βH226E mutations compromised TCR expression (*22*), and the βN225A/βH226A double mutation sharply suppressed IL-2 secretion in a T cell hybridoma (*26*). Cryo-EM studies further identified coupling between the Cα DE loop and the CD3δ AB and DE loops (*28*), with substitutions impairing surface assembly (*22*). For example, αR185 abolished calcium signaling and reduced CD69 and ERK phosphorylation (*24*), while αD183K mutation impaired CD69 and ICOS induction (*27*). Additionally, mutations in the CD3ε H strand and CD3γ AB loop decreased surface expression (*22*).

Beyond these previously mapped sites, our ΣOccupancy analysis uncovered new dynamic *cis*-contacts not captured in structural studies (*28-31*) yet known to affect TCR surface expression (*22*) and effector functions (*24, 33, 112*). Several of these align with mutagenesis phenotypes, such as the Cα AB loop interaction with the CD3ε FG loop, proposed to undergo conformational shifts during ligand-induced activation (*33, 112, 113*). Mutation αK148E, and substitutions within CD3ε FG loop prevented stable formation of TCR–CD3 assembly at the cell surface (*22*). Likewise, the Cα DE loop contacts CD3δ AB loop, where αM187 substitution impaired calcium signaling (*24*). Not all high-occupancy residues have been fully explored (fig. S6). For example, the Cβ G strand dynamically interacts with CD3ε and CD3γ; βS255R enhances TCRβ surface expression, whereas βI253A and βE257A mutations did not alter expression (*22*). However, βE257K and βW259Y mutations impaired IL-2 production and reduced CD69 expression (*27*). From an engineering perspective, these cross-validated and newly identified *cis*-interacting residues represent potential levers to modulate signaling outputs without perturbing antigen recognition, offering strategies to fine-tune T cell efficacy in immunotherapy.

Intriguingly, ΔOccupancy also revealed force-sensitive TCR–CD3 residues, highlighting their role in mechanotransduction. For example, the βS255 and βW259 contacts on the Cβ G strand are force-enhanced, yet the stabilizing βW259Y mutation paradoxically resulted in loss-of-function (*27*). This supports the view that certain force-strengthening TCR–CD3 ECD anchors (*36, 114*) act as potential "motion stoppers", particularly enriched at the membrane-proximal region of CD3δε. Alterations of these residues could tune CD3δε fluctuations under force, thereby enhancing or dampening activation. Furthermore, ΔOccupancy can be readily extended from force *vs.* no force to *apo vs. holo* conditions, providing a framework for generating new testable hypotheses on TCR triggering.

### SUPPLEMENTARY TABLES

**Table S1. Kinetic parameters in Fig. 1, C to E of the main text by the MAF and BFP assays.** Summary of effective 2D affinities, effective 2D on-rates, and 2D off-rates for the TCRαβ–CD3γε, TCRαβ–CD3δε, and CD3δε–CD3γε ECD (purified proteins) interactions, as well as 2B4 TCR (expressed on hybridoma cells) interaction with K5:I-$E^k$ pMHC (purified proteins). Data represent mean ± SEM.

| Interactions | Effective 2D affinity ($\mu m^4$) | Effective 2D on-rate ($\mu m^4\ s^{-1}$) | 2D off-rate ($s^{-1}$) |
|---|---|---|---|
| TCRαβ–CD3γε | $3.8 \pm 0.7 \times 10^{-6}$ | $5.3 \pm 1.0 \times 10^{-5}$ | 14.12 ± 0.15 |
| TCRαβ–CD3δε | $2.8 \pm 0.6 \times 10^{-6}$ | $3.1 \pm 0.7 \times 10^{-5}$ | 11.13 ± 0.48 |
| CD3γε–CD3δε | $1.1 \pm 4.4 \times 10^{-5}$ | $1.6 \pm 0.7 \times 10^{-4}$ | 14.89 ± 0.37 |
| 2B4–K5:I-Ek | $5.9 \pm 0.4\times 10^{-4}$ | $5.0 \pm 0.8 \times 10^{-3}$ | 8.48 ± 1.22 |

**Table S2. Kinetic parameters of catch-slip bond models in Fig. 2D of the main text.** Summary of the best-fit Bell parameters of two-state, two-pathway dissociation models for the TCRαβ–CD3γε (*top*), TCRαβ–CD3δε (*middle*), and CD3δε–TCRαβ–CD3γε (*bottom*) species, respectively.

| **Transition** | **Rate** | $k_{i,j}^0$ (1/s) | $x_{i,j}$ (Å) |
|---|---|---|---|
| TCRαβ–CD3γε → 0 | $k_{-f,\gamma}$ | 11.1 ± 0.298 | 0 ± 0.0685 |
| (TCRαβ–CD3γε)* → 0 | $k_{-s,\gamma}$ | 0.196 ± 0.0228 | 4.40 ± 0.304 |
| TCRαβ–CD3γε → (TCRαβ–CD3γε)* | $k_{a,\gamma}$ | 1.60 ± 0.355 | 1.93 ± 0.801 |
| TCRαβ–CD3γε ← (TCRαβ–CD3γε)* | $k_{-a,\gamma}$ | 15.0 ± 1.91 | - 32.5 ± 2.30 |
| **Transition** | **Rate** | $k_{i,j}^0$ (1/s) | $x_{i,j}$ (Å) |
| TCRαβ–CD3δε → 0 | $k_{-f,\delta}$ | 11.4 ± 0.162 | 0 ± 0.0652 |
| (TCRαβ–CD3δε)* → 0 | $k_{-s,\delta}$ | 0.184 ± 0.0412 | 2.80 ± 0.422 |
| TCRαβ–CD3δε → (TCRαβ–CD3δε)* | $k_{a,\delta}$ | 1.50 ± 0.316 | 0 ± 0.504 |
| TCRαβ–CD3δε ← (TCRαβ–CD3δε)* | $k_{-a,\delta}$ | 8.77 ± 2.49 | - 12.3 ± 1.57 |
| **Transition** | **Rate** | $k_{i,j}^0$ (1/s) | $x_{i,j}$ (Å) |
| CD3δε–TCRαβ–CD3γε → 0 | $k_{-f,\gamma\delta}$ | 0.700 ± 0.204 | 5.45 ± 0.204 |
| (CD3δε–TCRαβ–CD3γε)* → 0 | $k_{-s,\gamma\delta}$ | 0.0519 ± 0.0196 | 0 ± 0.859 |
| CD3δε–TCRαβ–CD3γε → (CD3δε–TCRαβ–CD3γε)* | $k_{a,\gamma\delta}$ | 1.00 ± 0.118 | 0 ± 0.322 |
| CD3δε–TCRαβ–CD3γε ← (CD3δε–TCRαβ–CD3γε)* | $k_{-a,\gamma\delta}$ | 21.8 ± 9.82 | - 26.7 ± 3.21 |

**Table S3. Summary of identified high-occupancy residues, cross-referenced with previous structural evidence and mutagenesis studies.** The critical contacts of the CD3δε–TCRαβ–CD3γε interfaces (as identified by high ΣOccupancy in fig. S6) are listed by their chain-, region-, and residue-IDs. Their relevance was cross-validated through a literature search. The references with structural evidence are enumerated for a particular residue, for which *cis*-contacts have been previously identified by NMR, cryo-EM, *etc*. The references for the mutagenesis studies are also listed, where contacting residues have been mutated before, leading to either surface-expression-level alterations of TCR–CD3s or functional variations of downstream signaling, such as $Ca^{2+}$ induction, IL-2 secretion, *etc*. The rows in *red* highlight the critical contacts that are also strengthened by force (as identified by high ΣOccupancy and ΔOccupancy in fig. S6).

| Chain | Region | Residue | Expression | Functional reference | Structural reference |
|---|---|---|---|---|---|
| TCRα | AB loop | K148 | -- | | |
| | DE loop | R185 | - | calcium flux reduction (*24*) | Cryo-EM (*28*) |
| | | S186 | --- | change in activation (*24*) | Cryo-EM (*28*) |
| | | M187 | -- | calcium flux reduction (*24*) | |
| TCRβ | CC' loop | N181 | --- | | Cryo-EM (*28*) |
| | | G182 | -- | IL-2 reduction (*26*) | |
| | Helix 4 | S216 | + | | NMR (*25*) |
| | | T218 | -- | | NMR (*25*) |
| | | F219 | --- | | NMR (*25, 26*) |
| | Helix 4–F strand | P223 | - | IL-2 increase (*26*) | |
| | | R224 | --- | | NMR (*25, 26*) |
| | | N225 | | IL-2 reduction (*26*) | NMR (*25, 26*) |
| | | H226 | -- | IL-2 reduction (*26*) | Cryo-EM (*28*) |
| | G strand | I253 | | | Cryo-EM (*28*), NMR (*25*) |
| | | S255 | + | IL-2 increase (*27*) | Crosslink (*27*) |
| | | E257 | | | Crosslink (*27*) |
| | | W259 | + | IL-2 increase (*27*) | Crosslink (*27*) |
| CD3δ | AB loop | L29 | --- | | Cryo-EM (*28*) |
| | | E30 | --- | | Cryo-EM (*28*) |
| | | D31 | --- | | |
| | | R32 | --- | | |
| | EF loop | K62 | -- | | Cryo-EM (*28*) |
| | | I64 | -- | | Cryo-EM (*28*) |
| CD3ε | CD loop | N62 | -- | | |
| | | D63 | - | | |
| | | K64 | - | | |
| | FG loop | L90 | - | | Cryo-EM (*28*) |
| | | E91 | - | | |
| | H strand | R115 | --- | impaired activation, IL-2 (*21*) | |
| | | (R117) | --- | impaired activation, IL-2 (*21*) | |
| CD3γ | AB loop | Y34 | + | | |
| | | D35 | | | Cryo-EM (*28*) |
| | | Y36 | --- | | Cryo-EM (*28*) |
| | | Q37 | -- | | |
| | | E38 | --- | | |
| | | G40 | --- | | |
